# Predictive neural processing in adult zebrafish depends on shank3b

**DOI:** 10.1101/546457

**Authors:** Kuo‐Hua Huang, Peter Rupprecht, Michael Schebesta, Fabrizio Serluca, Kyohei Kitamura, Tewis Bouwmeester, Rainer W. Friedrich

## Abstract

Intelligent behavior requires a comparison between the predicted and the actual consequences of behavioral actions. According to the theory of predictive processing, this comparison relies on a neuronal error signal that reflects the mismatch between an internal prediction and sensory input. Inappropriate error signals may generate pathological experiences in neuropsychiatric conditions. To examine the processing of sensorimotor prediction errors across different telencephalic brain areas we optically measured neuronal activity in head-fixed, adult zebrafish in a virtual reality. Brief perturbations of visuomotor feedback triggered distinct changes in swimming behavior and different neuronal responses. Neuronal activity reflecting sensorimotor mismatch, rather than sensory input or motor output alone, was prominent throughout multiple forebrain areas. This activity preceded and predicted the transition in motor behavior. Error signals were altered in specific forebrain regions by a mutation in the autism-related gene shank3b. Predictive processing is therefore a widespread phenomenon that may contribute to disease phenotypes.

## Introduction

Our perception of the world is determined not only by external stimuli but also by our internal expectations. For example, self-tickling is always less effective than tickling by another person because the brain predicts the consequences of self-generated behavior and modifies perception accordingly (Blakemore et al, 2000). The framework of predictive processing proposes that predictions are generated based on an internal model of the world that is constantly updated based on an individual’s experience (Mumford, 1991; Rao and Ballard, 1999; Friston, 2008; Keller and Mrsic Flogel 2018). The internal model is thought to generate primarily inhibitory neural activity that suppresses excitatory responses of projection neurons to predictable components of sensory inputs. The remaining excitatory activity contains information about the prediction error and is used to update the internal model. Information about the world is thus encoded in the functional organization of neural networks that are continuously adapted to minimize a distributed error signal. This mechanism allows for efficient information storage and for continuous interactions between sensory inputs and memories.

A key prediction of this framework is that unexpected sensory inputs elicit responses in specific subsets of neurons that represent a prediction error (Huang and Rao 2011; Keller and Mrsic Flogel 2018). Consistent with this assumption, neural activity related to prediction errors has been observed in the cerebellum (Blakemore and Wolpert, 1998) and in the dopaminergic reward system (Schultz et al., 1997). Moreover, neural error signals have been studied in sensorimotor systems by quantifying responses to unexpected changes in the coupling between self-generated movements and sensory feedback. These studies identified neurons in visual (Keller et al., 2012; Saleem et al., 2013, Fiser et al., 2016; Stanley and Miall, 2007) and auditory cortices (Eliades and Wang, 2008; Keller and Hahnloser, 2009) that are driven by a mismatch between the actual and the predicted sensory input, rather than by sensory input or motor output alone. Neuronal circuits in sensory cortices may therefore function as comparators that compute the mismatch between the actual sensory input and a prediction generated by an internal model based on motor commands (Keller and Mrsic Flogel 2018). Moreover, it has been proposed that predictive processing is not limited to sensorimotor systems but of broad relevance also in other domains including cognitive processes that infer the mental states and actions of other individuals (Koster-Hale and Saxe, 2013). However, further studies are required to analyze predictive processing in different species, brain areas and tasks.

In the framework of predictive processing it has been suggested that pathological mental experiences are the consequence of an imbalance between internally generated predictions and stimulus-driven external inputs. Positive symptoms of schizophrenia including hallucinations may be generated by an excessive internal prediction that is not matched by an external sensory input of appropriate strength (Corlett et al., 2009; Fletcher and Frith, 2009; Frith, 2000). Conversely, symptoms of autism may be due to a “sensory overload” that occurs when responses to external sensory inputs are not matched by sufficiently strong and specific internal predictions (Pellicano and Burr, 2012; Friston et al., 2013). This scenario suggests that stereotyped and repetitive behaviors, which are hallmarks of autism spectrum disorders, represent a simple strategy to reduce sensory overload by making sensory input more predictable (Lawson et al., 2014; Lawson et al., 2017; Sinha et al., 2014).

Mechanistic analyses of predictive processing would benefit from a small animal model that allows for the systematic analysis of neural activity across multiple brain areas. In adult zebrafish, neural activity can, in principle, be measured by multiphoton calcium imaging in dorsal forebrain areas that are thought to be homologous to the basolateral amygdala (area Dm), isocortex (Dc) and possibly hippocampus (Dl) (Lal et al., 2018; Aoki et al., 2013; Mueller 2011). Moreover, in contrast to zebrafish larvae, adult zebrafish exhibit pronounced social behavior (Buske and Gerlai 2011; Laan et al., 2018), robust learning in associative and operant paradigms (Valente et al., 2012; Namekawa et. al, 2018; Doyle et al, 2017), and other complex behaviors. Adult zebrafish may therefore be an interesting model system to examine predictive processing in the context of different behaviors and genetic disease models.

Predictive processing can be analyzed in a virtual reality (VR) that offers the opportunity to generate specific mismatches between self-generated behavior and sensory feedback during simultaneous measurements of neural activity and behavior (Portugues and Engert 2011; Ahrens et al., 2012; Keller et al., 2012). Although simple virtual reality systems have been developed for zebrafish larvae (Ahrens et al., 2013), this approach has not yet been established in adult zebrafish. We therefore developed procedures for multiphoton calcium imaging of neural activity in the forebrain of head-fixed adult zebrafish during behavior in a high-resolution virtual reality. Perturbations of visuomotor coupling revealed a substantial fraction of forebrain neurons that responded specifically to visuomotor mismatch and predicted subsequent behavioral responses. The occurrence of these mismatch-sensitive neurons was increased in Dm and Dc but decreased in Dl of zebrafish carrying a mutation in shank3b, a gene that has been causally linked to an autism spectrum disorder in humans (Durand et al., 2007; Leblond et al, 2014). Predictive processing therefore occurs in multiple forebrain areas of a teleost, indicating that it is an important neural computation also in non-mammalian species. Moreover, these results are consistent with the hypothesis that symptoms of autism spectrum disorders are due to aberrant prediction error signals in specific brain areas.

## Results

### Head-fixation and Virtual Reality for Adult Zebrafish

To identify anchor points for the stable head-fixation of adult zebrafish we examined the skull after removing soft tissues. The adult skull consists of five major sets of bone: two sets cover the top of the forebrain and rostral midbrain, and another two sets cover the midbrain and the cerebellum. These parts of the skull are thin (<100 μm) and provide only limited mechanical support. The fifth set extends from the nostrils to the spinal cord and surrounds the brain ventrally and laterally (Figure 1A and 1B). This solid piece of bone is covered by muscle and connective tissues except for a site lateral to the cerebellum where the bone is located immediately underneath the skin. We took advantage of this location to glue an L-shaped head bar onto the bone after removal of the skin. The head bar was made of stainless steel and attached to the bone using tissue glue and dental cement (Figure 1C and Figure S1, also see Methods). The anchored bone encased and stabilized the entire brain, providing sufficient stability for in vivo imaging of neuronal activity at single-neuron resolution even during vigorous swimming.

**Figure 1:**
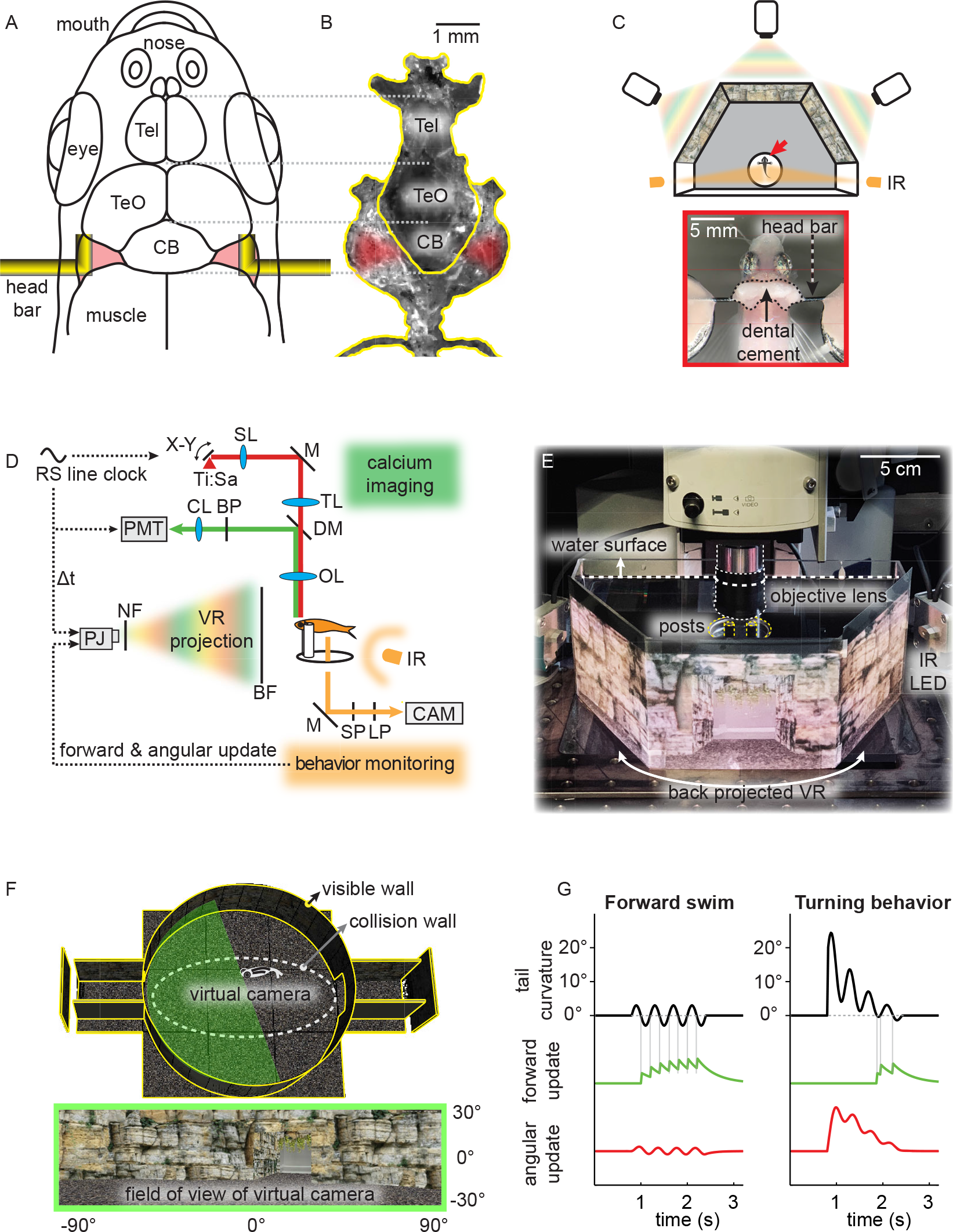
Head-fixation of Adult Fish, Closed-loop Virtual Reality (VR) and Neuronal Calcium Imaging during Behavior. (A) Attachment sites of L-shaped bars for head fixation (see also Figure S1). Tel: telencephalon; TeO: optic tectum; CB: cerebellum. (B) Skull bones corresponding to the drawing in A. (C) Top: schematic of VR projected by three projectors onto a panoramic screen with a 180 degree field of view. Arrow depicts head-fixed fish. Bottom: dorsal view of head-fixed fish. (D) Synchronization of closed-looped VR and two-photon imaging. The photomultiplier tube (PMT) and projector LEDs are gated in a non-overlapping manner by the line clock of the resonant scanner (RS). Ti:Sa: titanium-sapphire laser; SL: scan lens; M: mirror; TL: tube lens; DM: dichroic mirror; BP: band pass filter; PJ: projector; NF: notch filter; SP: short pass filter; LP: long pass filter; CAM: camera; BF: back-projection film. (E) Back-projection of VR onto a semi-hexagonal tank under the microscope. (F) 3D model of the VR consisting of a cylindrical arena, a flat floor and two tunnels (top). Walls were textured with naturalistic rocks and plants (bottom). A set of three virtual cameras together capture a 180 degree field of view (green shade). An invisible collision wall (dashed line) kept virtual cameras away from the VR boundaries to prevent pixelation of the texture. (G) Algorithm to update the virtual cameras based on tail movements of head-fixed animals. Forward movement in the VR (green) is triggered by the zero crossings of the caudal tail curvature (black). Angular movement in the VR (red) is proportional to the caudal tail curvature after low-pass filtering.

To enable locomotory behavior under head fixation we developed a closed-loop VR setup that allowed head-fixed animals to move in a horizontal plane within a 3D virtual environment (Figure 1C, 1F and Figure S2A). The VR had an update rate of 50 Hz and was projected onto the walls of a custom-made tank from the backside, covering a 180-degree field of view. The use of three projectors allowed us to generate VR textures with high resolution and contrast (Figure 1E). Fish were illuminated laterally by an infrared light source and the tail was filmed continuously with a video camera at 50 Hz.

To couple swimming behavior to displacements in the VR we predicted swim trajectories from the curvature of the tail in real time. Symmetric tail movements resulted in the forward translation of virtual cameras in the VR (forward swimming). This mapping was implemented by triggering translations on the zero-crossings of the caudal tail curvature, which capture the back strokes of the tail movement that propel the fish forward during free swimming. Asymmetric tail undulations resulted in an initial change in heading direction, followed by a brief forward movement (Figure 1G).

Without head fixation, adult zebrafish typically swim for extended periods of time in discrete bouts. In the home tank, these swim events often occur at quasi-periodic intervals and consist of one or multiple tail flicks followed by a short gliding period. We observed that head-fixed adult zebrafish in a VR without closed-loop sensorimotor coupling struggled vigorously, presumably reflecting a high level of anxiety, and subsequently entered a prolonged period of inactivity. Under closed-loop conditions, in contrast, animals typically performed spontaneous quasi-periodic swim events. Each swim event consisted of one or multiple tail undulations followed by a short pause before the onset of the next event. The mode of the duration of swim events was 0.66 s (n = 37 fish, Figure S2B). This swimming behavior was very similar to the swimming pattern of freely behaving animals. Naturalistic swimming behavior under closed-loop conditions was observed for more than 6 hours (Figure S2C). These observations indicate that head-fixed animals in a VR with appropriate sensorimotor feedback did not experience high levels of stress.

### Neuronal Activity in the Dorsal Telencephalon of Adult Zebrafish

To perform two-photon calcium imaging during behavior in the VR we temporally segregated photons of the VR from fluorescence measurements. This was achieved by exclusive gating of the projectors’ LEDs and the photomultiplier tube using the line clock of the 8 kHz resonance scanner as a trigger signal (Figure 1D; Methods). Calcium signals were measured in neuroD:GCaMP6f transgenic fish that expressed the calcium indicator GCaMP6f in most pallial forebrain areas at adult stages (Rupprecht et al., 2016). Because the bones over the dorsal telencephalon are very thin (~50 μm), multiphoton calcium imaging could be performed through the intact skull up to 200 µm below the brain surface (Figure 2A & 2B, also Figure S3). The maximal dimensions of the adult telencephalon are approximately 1 mm x 1.2 mm x 0.6 mm (length x width x height). The optically accessible volume includes a large portion of this volume and comprises multiple brain areas in the dorsal pallium. We focused on telencephalic areas Dm (medial zone of the dorsal telencephalon), which is likely to be homologous to the basolateral amygdala and related areas (Lal et al., 2018), the dorsorostral and dorsocaudal parts of area Dc (rDc and cDc, respectively; central zone of the dorsal telencephalon), which are thought to be homologous to isocortex (Mueller et al., 2011; Aoki et al, 2013), and the dorsal part of Dl (lateral zone of the dorsal telencephalon). The relationship of Dl to the mammalian brain is not fully understood but the ventral part of Dl is likely to be related to the hippocampus (Rodriguez et. al., 2002).

**Figure 2:**
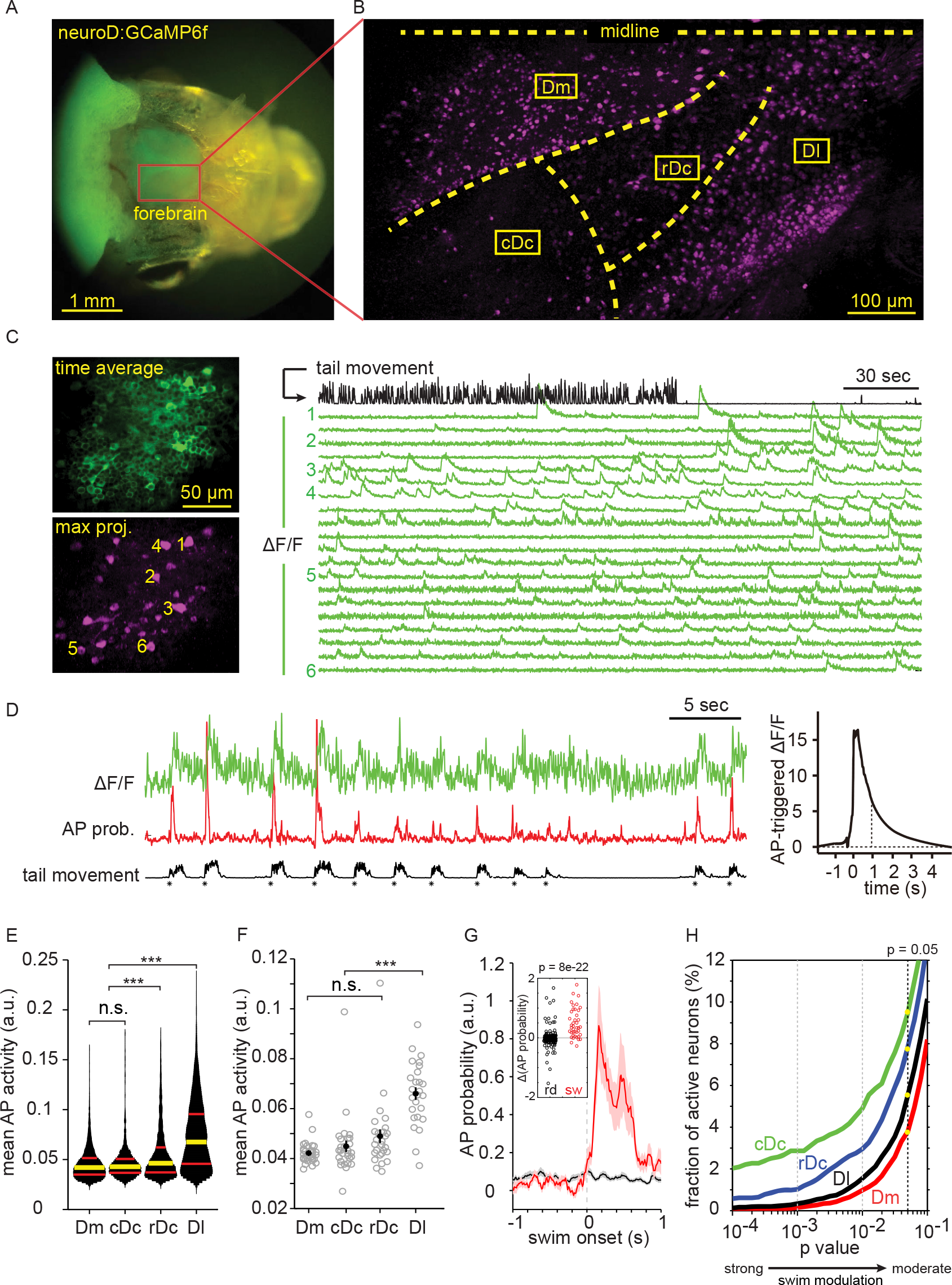
Region-specific Neuronal Activity in the Telencephalon of Adult Zebrafish. (A) Head of an adult zebrafish expressing neuroD:GCaMP6f, imaged through a wide-field epifluorescence microscope. (B) Neuronal activity pattern of adult zebrafish forebrain imaged using a two-photon microscope through the intact skull after skin removal (also see Figure S3). Active neurons (magenta) were revealed by a maximum projection of pixel intensities along the time axis of a mean-subtracted image series (90 s). (C) Simultaneous recording of neuronal activity and behavior. Upper left: time-averaged raw fluorescence. Lower left: maximum intensity projection of mean-subtracted image series, revealing active neurons. Right: traces of calcium signals (ΔF/F) from individual neurons (green; numbers indicate ROIs in the maximal projection image) together with the simultaneously recorded tail movement (black). (D) Calcium signals of a swim-modulated cell (ΔF/F; green), estimated AP probability (red), and intensity of tail movement (black). Asterisks depict onsets of swim events. Right: AP-triggered average calcium transient. Dashed line depicts decay time constant of 0.9 s. (E) Fraction of neurons as a function of activity (AP probability) in the dorsal forebrain during spontaneous behavior in a closed loop VR. Each neuron was recorded for 267 s. Yellow and red bars indicate median and interquartile range, respectively (Dm: n = 7449 neurons; cDc: n = 1675 neurons; rDc: n = 2941 neurons; Dl: n = 7967 neurons from 32 animals). (F) Mean AP probability in different brain regions and animals. Each circle represents an animal (n = 31, 28, 27, 26 fish for Dm, cDc, rDc, Dl, respectively; error bar represents SEM). Neural activity was significantly higher in Dl than in other brain areas (one way ANOVA followed by Tukey’s HSD procedure for multiple comparisons; *** = adjusted p value < 0.001). (G) Mean AP probability triggered on swim onset (red) and on random time points (black) of a swim-modulated neuron (shading represents SEM). Inset shows the difference in AP probability between 0.5 s time windows immediately before and after the onset of swim events (n = 126 events). A neuron is considered swim-modulated when the distribution of swim-triggered responses is significantly different (p < 0.05; t-test) from randomly-triggered responses. (H) Fraction of neurons that were classified as swim-modulated as a function of detection stringency (p-value). Dashed line (black) depicts the fraction of swim-modulated neurons for p < 0.05 in cDc (9.4%), rDc (7.7%), Dl (5.5%) and Dm (3.8%).

During spontaneous swimming in a closed-loop VR, individual neurons showed sparse and discrete fluorescence transients. To highlight neurons that were active within a given time window we performed a maximum projection of pixel intensities along the time axis of an image series and subtracted the mean image. In the resulting difference image, bright pixels thus represented somata of neurons that showed at least one calcium transient (Figure 2C). Because the dynamics of somatic calcium transients is relatively slow (Figure 2D, decay time constant 0.9 s) in comparison to behavioral variables, we estimated the underlying fluctuations in action potential (AP) probability using a convolutional neural network (Berens et al., 2018; Figure 2D). During spontaneous swimming, we determined the mean AP probability of individual neurons during a time window of 267 seconds. We found that spontaneous activity was significantly higher in Dl and rDc in comparison to Dm and cDc (median = 0.07, 0.05, 0.04, 0.04 a.u., n = 7967, 2941, 7449, 1675 cells, respectively; Kruskal-Wallis test followed by Tukey’s HSD for multiple comparisons; Figure 2E). The elevated level of neural activity in Dl was also observed when mean activity levels were compared across animals (n = 31 fish, Figure 2F). This analysis showed region-specific neural activity during spontaneous motor behavior in the dorsal telencephalon.

We next analyzed the modulation of neuronal activity during self-generated motor behavior in the virtual reality. In a subset of neurons, fluctuations in calcium signals and AP probability were obviously correlated to discrete swim events (Figure 2D). To analyze this relationship more systematically we selected the onsets of swim events as trigger points for the analysis of AP probability and calculated the difference in AP probability Δp between 0.5 s time windows immediately before and after the trigger for each neuron. As a control, we used randomly chosen time points as triggers. Neurons were considered swim-modulated when the distribution of Δp was significantly different between onset-triggered and randomly triggered conditions (one-sided t-test with p < 0.05, Figure 2G). We found that swim-modulated neurons were more prevalent in the Dc regions (cDc: 9.4%, 158/1675 cells; rDc: 7.9%, 233/2941 cells) than in Dl (5.5%, 438/7967 cells) and Dm (3.7%, 275/7449 cells). This region-specific difference in swim modulation remained qualitatively similar and became quantitatively more distinct when the stringency of the criterion to define swim modulation was increased (Figure 2H). The dorsal part of Dc may therefore include motor-related areas or receive strong inputs from such areas.

### Behavioral Responses to a Visual Feedback Mismatch

To generate brief episodes of visuomotor mismatch, we introduced a left-right reversal to the angular update of the VR for 5 – 10 s (mean duration 7.5 s, Figure 3A, also see Movie S1). These visuomotor perturbations were generated every 2 – 7 minutes with a mean inter-event interval of 6 minutes. Visuomotor perturbations did not affect the linear update of the VR (forward swimming) but resulted in an unexpected rotation of the VR in the direction opposite to the animal’s expectation during turns. Because the VR was updated based on the tail movement, the animal experienced the left-right reversal only during episodes of active swimming.

**Figure 3:**
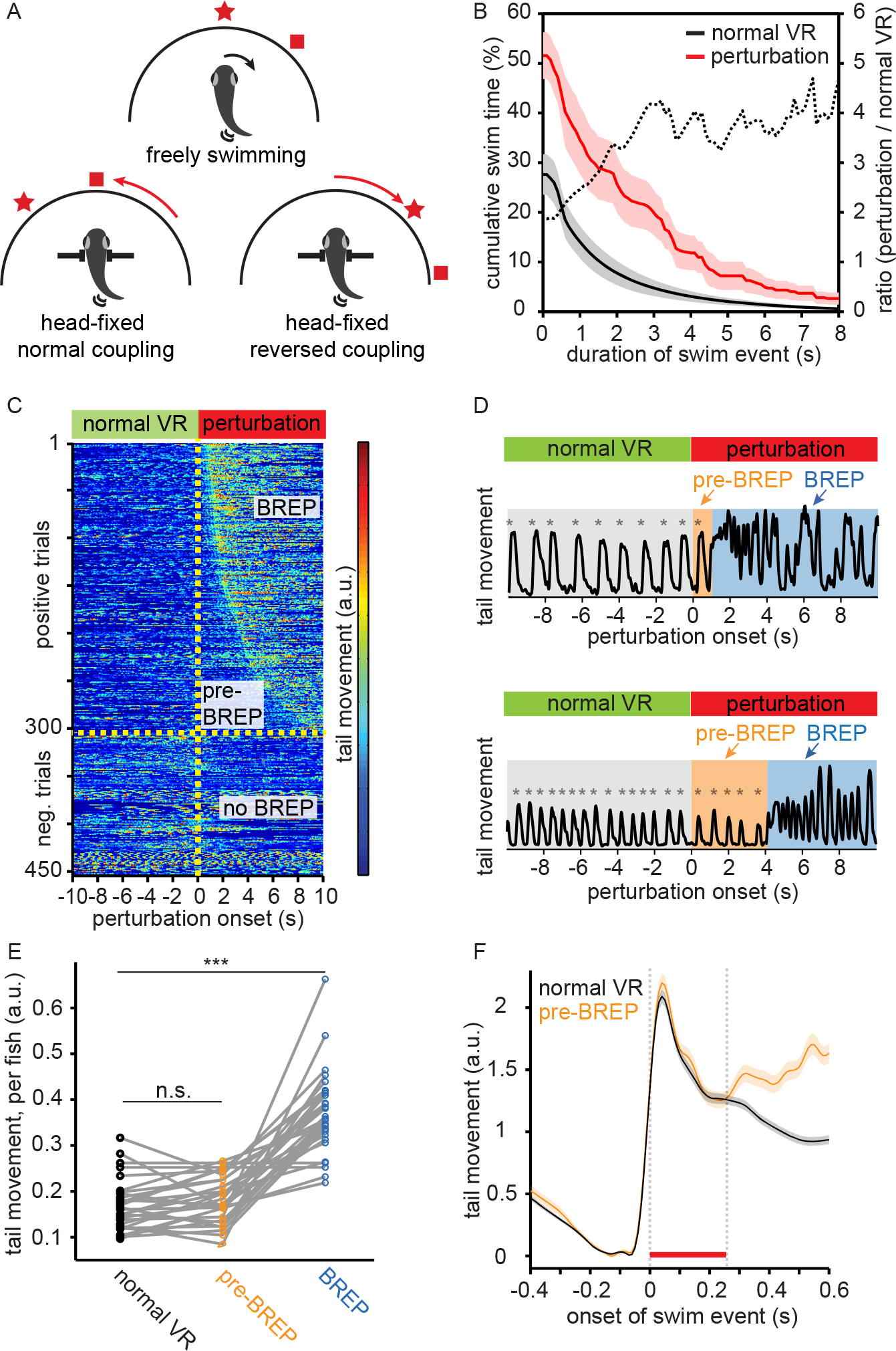
Behavioral Responses to a Visual Feedback Mismatch. (A) Schematic showing VR updates during normal and reversed visuomotor coupling. (B) Cumulative distribution of time engaged in swim events of different duration. Swimming activity was elevated during visuomotor perturbations (red) in comparison to swims during normal visuomotor coupling (black). The fraction of long swimming events was also increased during perturbations (dashed line). Shading shows SEM over animals (n = 32). (C) Tail movement (color-coded) in different perturbation trials (rows) as a function of time (x-axis). A visuomotor perturbation of 10 s duration was introduced at t = 0. Trials from 32 fish were sorted by the onset of the behavioral response (BREP). In 33% of the trials, no BREP was observed (bottom). In the remaining trials, the delay varied over a wide range. The intensity of tail movements was quantified by the mean of absolute pixel-wise difference in successive video frames. (D) Examples of tail movement before the VR perturbation (normal VR), during perturbation but before behavioral responses (pre-BREP period), and during behavioral responses (BREP period). Asterisks indicate the onset of swim events. Top: short delay (trial 58 in C). Bottom: longer delay (trial 240 in C). Intensity of tail movement was quantified as in C. (E) Intensity of tail movement (quantified as in C) during a normal visuomotor coupling, during the pre-BREP period and during the BREP period. Each plot symbol represents one fish (normal VR vs. pre-BREP, p = 0.07, pairwise t-test; normal VR vs. BREP, p = 1.0e-9, pairwise t-test; n = 32 fish). (F) Mean intensity of tail movement (quantified as in C) triggered on the onset of swim events during normal visuomotor coupling and during the pre-BREP period. The initial structure of swim events is indistinguishable between the two conditions in the first 230 ms (red bar). Swim events within a perturbation trial was averaged. Mean and SEM were calculated over trials (n = 303 trials).

To analyze the behavioral response, we quantified the intensity of tail movements by computing the mean of absolute pixel-wise difference in adjacent frames of behavioral recordings. We found that the fraction of time that fish were engaged in active swimming increased from 26% during normal visuomotor coupling to 51% during left-right reversals (Figure 3B). In addition, the fraction of time that fish performed long swim events (> 2 s) increased from 8% during normal visuomotor coupling to 26% during left-right reversal (Figure 3B). Hence, both the overall swimming time and the duration of swim events increased during the perturbation.

Detailed visual inspection of videos (50 Hz, 450 perturbation trials) revealed that the perturbation of the VR usually triggered a distinct sequence of motor outputs that followed the onset of the perturbation with a variable delay (Movie S2). The sequence typically started with 1 – 5 unilateral tail bends of increasing frequency and amplitude at the caudal part of the tail. These tail movements therefore resembled J-turns, which are orienting movements that turn the body axis during prey capture (McElligott & O’Malley, 2005; Bianco et al., 2011). Initial J-turn-like movements were then followed by vigorous and irregular movements of the entire tail. We used these features to define a behavioral response to perturbation (BREP) and manually annotated BREPs in each trial. Trials were then sorted by the time of BREP onset, which was defined as the first noticeable change in motor output (onset of the J-turn-like movements). Visual analysis of tail movement intensity across all trials confirmed that BREP onset represented a distinct transition in behavioral state from normal swimming to prolonged and vigorous tail movements (Figure 3C). Based on these observations we divided the time following the perturbation into two phases in each trial: the pre-BREP period corresponded to the delay period between the perturbation and the BREP, and the BREP period corresponded to the time between BREP onset and the end of the 10 s analysis time window.

The duration of the pre-BREP period varied substantially between trials (median = 2.1 s, interquartile range 1.2 – 4.1 s). During this time, animals continued to perform short periodic swims similar to those before the perturbation (Figure 3D). The intensity of tail movement was not significantly different between swims during normal visuomotor coupling and during the pre-BREP period (Figure 3E, pairwise t-test, p = 0.07) but significantly higher during BREP period (pairwise t-test against normal coupling, p = 1.0e-9, n = 32 animals). Time-resolved analyses of swim-triggered tail movements further showed that the average tail movement intensities during normal coupling and during the pre-BREP period were virtually identical for the first 230 ms of a swim event (Figure 3F, red bar). Subsequently, the intensity of tail movements triggered during the pre-BREP period increased because some events included transitions to BREPs. These results show that the intensity and the temporal structure of short periodic swims were maintained during the pre-BREP period when the fish experienced the visual perturbation but did not yet undergo a transition in motor output.

### Neuronal Activity during Visual Feedback Mismatch

To investigate neuronal activity during VR perturbations we performed multiphoton calcium imaging during behavior. Each optical plane was scanned for 30 minutes while 1 – 7 brief perturbations (left-right reversals; 5 – 10 s per perturbation, mean duration 7.5 s) were introduced. In each animal, 3 – 6 different locations in the forebrain were examined in Dm, cDc, rDc and Dl. In most fish, activity was measured in all four brain areas. Among 7090 cells recorded from 32 animals, 882 cells (12.2%) responded to the VR perturbation (Dm 9.2%, 233/2520 cells; cDc 15.0%, 121/805 cells, rDc 19.4%, 171/882 cells, Dl 12.4%, 357/2883 cells; see Methods for the definition of responsive cells).

Responses to the VR perturbation may reflect multiple processes including (1) increased motor activity, (2) a change in emotional state, (3) preparatory activity before the behavioral transition, (4) the detection of the mismatch between the motor output and the visual feedback. As swimming did not change significantly during the delay between the perturbation and the BREP (ref. Figure 3E & 3F), responses during the pre-BREP period are unlikely to be motor-related. We therefore separated the analysis of responses during the pre-BREP and the BREP by aligning activity measurements on the onset of the BREP in each trial. We then clustered neurons based on their activity by affinity propagation (Frey and Dueck 2007) and found that in 5 out of 11 clusters a significant fraction of neurons increased their average activity already before the onset of the BREP (Figure 4A). Hence, a substantial fraction of neuronal responses did not directly reflect a change in motor output.

**Figure 4:**
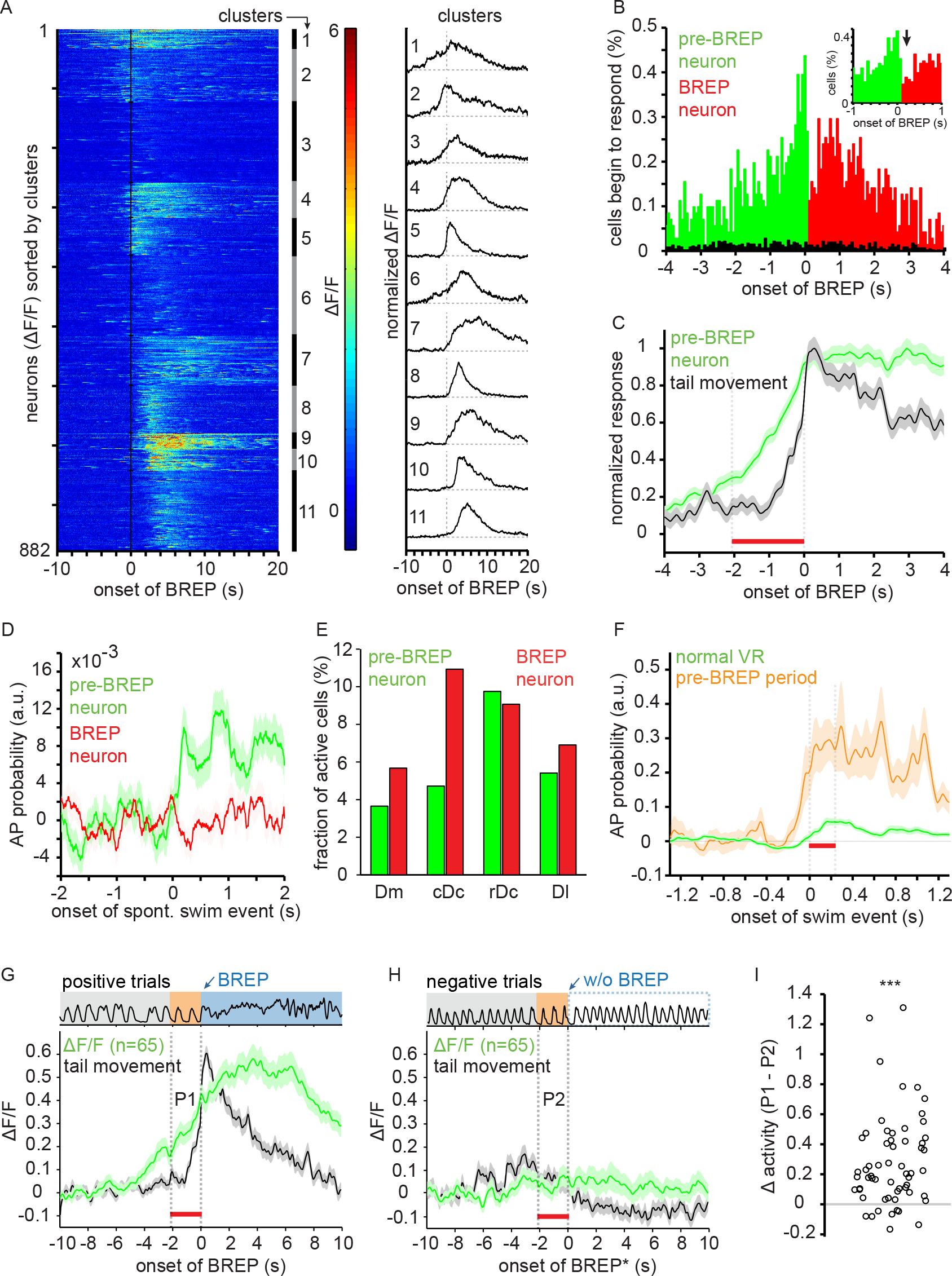
Visuomotor Mismatch Signal in the Zebrafish Forebrain. (A) Clustering of neurons by activity traces. Calcium signals (ΔF/F) of all responsive neurons (882 out of 7090 cells from 32 fish) were aligned on the onset of the BREP and clustered by affinity propagation. Right: average ΔF/F of each neuronal cluster. (B) Distribution of responsive neurons as a function of response onset relative to BREP onset. Green: neurons that started to respond after perturbation onset but before BREP onset (n = 382); Red: neurons that started to respond after BREP onset (n = 500). Black: fraction of active neurons detected around the onsets of spontaneous strong swims in the normal VR. The higher fraction of responsive neurons during visuomotor perturbations (red/green) indicates that most responses were triggered by visuomotor mismatch. (C) Time course of average pre-BREP neuronal activity (green) and tail movement intensity (black), normalized and aligned to BREP onset. Red bar indicates the median delay (2.06 s) from perturbation onset to BREP onset. (D) AP probability of pre-BREP neurons (green; n = 382) and BREP neurons (red; n = 500) triggered on the onset of spontaneous swims in a closed loop VR. (E) Prevalence of BREP-related neurons in the dorsal forebrain. Pre-BREP neurons: Dm, 3.7% (92 cells); cDc, 4.7% (38 cells); rDc, 9.8% (86 cells); Dl, 5.4% (156 cells). BREP neurons: Dm, 5.7% (143 cells); cDc, 10.9% (88 cells); rDc, 9.1% (80 cells); Dl, 6.9% (199 cells). Active neurons identified in Dm, cDc, rDc and Dl are 2520, 805, 882, 2883 cells, respectively (n = 32 fish). (F) AP probability of swim modulated pre-BREP neurons (n = 63) triggered on swim onset during normal visuomotor coupling (green) and during the pre-BREP period (orange). Red bar indicates the time period (230 ms) during which the intensity of the tail movement is indistinguishable between the two conditions (Figure 3F). (G, H) The same set of pre-BREP neurons show high average activity in positive trials (G, 124 trials) and no activity in negative trials (H, 106 trials). Red bar indicates the median delay (2.06 s) between perturbation onset and BREP onset in the positive trials. n = 65 neurons from 13 animals. (I) Activity difference between the pre-BREP periods during positive (P1 in G) and negative trials (P2 in H). Each symbol represents a neuron (n = 65). The activity of pre-BREP neurons during the pre-BREP period predicts the subsequent behavioral response (p = 4.1e-7, pairwise t-test).

Further analyses across all responsive neurons (n = 882) showed that response onset times of individual neurons were broadly distributed (Figure 4B). The probability of response onsets changed abruptly at BREP onset (Figure 4B, inset), suggesting that different functional subpopulations of neurons responded during the pre-BREP phase and during the BREP. We therefore divided responsive neurons into a subpopulation that started to respond during the pre-BREP period (pre-BREP neurons, n = 382) and a subpopulation that started to respond after the transition in motor behavior (BREP neurons, n = 500). The average activity of pre-BREP neurons increased well before the increase in motor output and remained elevated during the BREP (Figure 4C).

The activity of the pre-BREP and BREP neurons may be related primarily to the perturbation of visuomotor coupling or to a change in motor output. To distinguish between these possibilities we identified swim events similar to BREPs during normal visuomotor coupling (“spontaneous strong swims”; swim duration >2 seconds, Figure 3B) and quantified the neuronal activity during these events. The fraction of neurons that became active around the onset of these strong swims (737/64600 neurons; 1.1%, see Methods) was much lower than the fraction of neurons that responded to VR perturbations (882/7090 neurons; 12.2%). In addition, the distribution of response onset times was nearly uniform without a detectable discontinuity after the onset of the swim (Figure 4B, black). These results indicate that most pre-BREP and BREP responses were driven, directly or indirectly, by the VR perturbation. Moreover, these results confirm that the measured calcium signals do not represent motion artifacts caused by transitions in motor output.

To examine whether pre-BREP and BREP neurons showed different activity patterns during spontaneous behaviors in the normal VR we performed a swim-triggered average of their AP probability. Using the same criterion as before to define swim modulation (Figure 2G), we found that 16.5% (63/382) of the pre-BREP neurons but only 2.8% (14/500) of BREP neurons were significantly modulated by swimming. Hence, in pre-BREP neurons, but not in BREP neurons, the probability of swim-modulation is substantially higher than in the average population of neurons (5.6%, ref. Figure 2H). Consistent with this observation, the mean activity of pre-BREP neurons, but not BREP neurons, increased sharply at the onset of swimming (Figure 4D). These results support the notion that pre-BREP neurons and BREP neurons include functionally different neuronal populations.

Pre-BREP and BREP neurons were found in all telencephalic areas examined. The relative abundance of the two neuronal populations across brain areas were similar except that BREP neurons were more abundant in cDc (Figure 4E). In general, the abundance of BREP neurons across brain areas resembled the abundance of swim-modulated neurons (cDc > rDc > Dl > Dm; Figure 2H). Together, these results demonstrate that a substantial fraction of neurons in multiple telencephalic brain areas respond to a perturbation of sensorimotor feedback. These responses are unlikely to be directly involved in the preparation or control of motor behavior because they did not occur during spontaneous strong swims. BREP-related neural responses are therefore likely to represent sensorimotor mismatch signals.

### Visuomotor Prediction Error Signals in the Zebrafish Forebrain

The average activity of pre-BREP neurons was modulated by swims during normal visuomotor coupling, which may indicate that these neurons are directly involved in the control of motor behavior. Alternatively, pre-BREP neurons may represent an error signal that is generated by a comparison between sensory input and a motor-related prediction. In this scenario, swim-locked activity of pre-BREP neurons during spontaneous behavior may reflect a residual visuomotor mismatch in the VR. To distinguish between these possibilities, we analyzed the activity of the swim-modulated pre-BREP neurons (n = 63 cells) and compared their swim-triggered activity during two conditions: (1) spontaneous swims during normal visuomotor coupling, and (2) swims during the pre-BREP period, i.e. after onset of the visuomotor perturbation but prior to the transition in the behavioral state. For each neuron, a swim-triggered average of the AP probability was calculated. The average activity of these pre-BREP neurons during spontaneous swims showed a rhythmic pattern with a period consistent with the periodic swim events (mode = 0.66 s, Figure S2B). Interestingly, the swim-triggered activity of these neurons was strongly increased during the pre-BREP period (Figure 4F), even when the intensity and the structure of the motor outputs were indistinguishable from spontaneous swims (Figure 3F). The activity of these pre-BREP neurons therefore did not simply reflect the intensity of the motor output performed by the animal. Rather, it is likely that the phase-locked activity of pre-BREP neurons signaled an unexpected visual feedback during active swimming, indicating that pre-BREP neurons detect visuomotor mismatch.

Since the detection of a sensorimotor mismatch was required for the animal to respond to the perturbation, we expected that the prediction error signal should be weaker or absent in trials when the perturbation failed to elicit a behavioral response. To test this hypothesis we separated the perturbation trials into two groups: trials when a behavioral response occurred (positive trials) and trials without behavioral response (negative trials, ref. Figure 3C). This analysis was restricted to neurons that showed pre-BREP responses in at least 50% of the positive trials, and that were also recorded in negative trials (n = 65 neurons). In positive trials, we observed a sharp increase in tail movement intensity at BREP onset while the mean activity of pre-BREP neurons increased more gradually. The onset of this ramping activity preceded the increase in the tail movement intensity (Figure 4G). In negative trials, we used the median delay of BREPs in the positive trials to align the neuronal activity for a comparable analysis because animals did not show a BREP. We found that the same neurons showed no detectable response to the perturbation (Figure 4H). On average, the activity across the same pre-BREP neurons after onset of the perturbation was significantly higher in positive trials than in negative trials (Figure 4I, pairwise t-test, p = 4.1e-7). Hence, the activity of pre-BREP neurons predicted the subsequent behavioral response.

### Perturbation of Visuomotor Mismatch Signals by a Mutation in Shank3b

The hypothesis that the perceptual experience of autistic individuals may be a consequence of an impaired balance between sensory input and internal predictions prompted us to examine zebrafish with mutations in shank3, a scaffolding protein in the postsynaptic density of glutamatergic synapses (Naisbitt et al, 1999; Tu et al., 1999). Mutations in human shank3 cause Phelan-McDermid Syndrome, a disorder of the autism spectrum with high penetrance (Phelan and McDermid, 2011). Major symptoms of autism in humans are repetitive behaviors and impaired social interactions. In mice, a mutation in shank3 causes similar symptoms including self-injurious repetitive grooming and deficits in social interactions (Bozdagi et al., 2010; Peça et al, 2011; Jiang and Ehlers, 2013; Monteiro and Feng, 2017). The zebrafish genome contains two orthologs of human shank3, shank3a and shank3b, as a result of a recent genome duplication (Postlethwait et al., 2000; Kozol et al., 2015). Here we focused on the adult zebrafish with a mutation in shank3b. A disruption of this gene has been reported to result in repetitive behavior and reduced social interactions in adult zebrafish (Liu et al., 2018), consistent with phenotypes of shank3 mutations in other species. All comparisons were made between homozygous mutant and wild-type (wt) siblings from the same crossings.

Shank3b mutants (shank3b^−/−^) showed no obvious developmental or behavioral deficits upon superficial inspection as adults, consistent with previous observations (Liu et al., 2018). To confirm that shank3b mutants nevertheless showed altered responses to sensory input we first analyzed the behavior of freely swimming adult zebrafish towards a movie of conspecifics (Figure 5A). In this simple assay we quantified the fraction of time that fish spent near the movie screen during a 20 s presentation period. We found that shank3b^−/−^ fish spent significantly more time near the stimulus (55.4 ± 8.4%, n = 16) than wt siblings (31.4 ± 6.0%, n = 22 in wt siblings; t-test, p = 0.023; Figure 5B). The swim velocity and swim bout frequency were slightly but not significantly lower in the mutant (2.67 ± 0.08 cm/s and 0.25 Hz) as compared to wt siblings (2.82 ± 0.09 cm/s and 0.30 Hz; Figure S4A). The increased time spent near the movie can therefore not be explained by lethargic behavior but may reflect repetitive behavior towards the visual stimulus. To confirm that the observed phenotype can be attributed to a loss of function of shank3b we further examined transgenic animals expressing a dominant negative isoform of shank3b under a broadly expressing promoter (elavl3:shank3b^ΔC^). As observed in shank3b^−/−^ fish, elavl3:shank3b^ΔC^ transgenics spent a significantly higher fraction of time near the movie (75.1 ± 2.8%, n = 10) when compared to wt siblings (43.6 ± 8.9%, n = 14; t-test, p = 0.008). The average swim velocity and swim bout frequency (2.58 ± 0.15 cm/s and 0.26 Hz) were again slightly but not significantly lower than in wt siblings (2.63 ± 0.20 cm/s and 0.35 Hz; Figure S4B). Hence, shank3b mutants and dominant-negative transgenics showed consistent and specific behavioral phenotypes, indicating that these phenotypes are the consequence of a loss of shank3b function.

**Figure 5:**
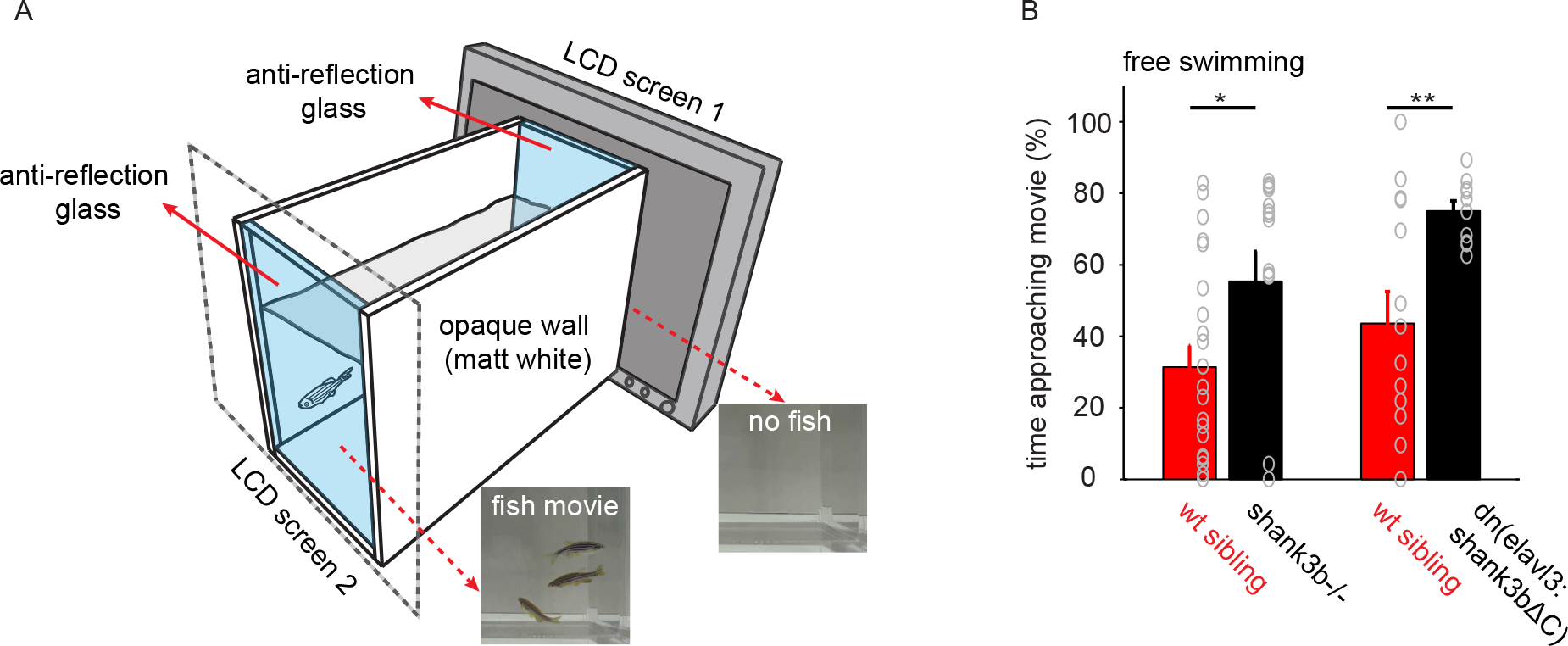
Loss of Shank3b Function Results in Consistent Behavioral Phenotype. (A) Schematic of the setup to test behavioral responses of freely swimming fish to movies of conspecifics. (B) Left: Shank3b^−/−^ fish spent significantly more time near a 20 s movie presentation of conspecifics than wt siblings (shank3b^−/−^: 55.4 ± 8.4%, n = 16 fish. Wt siblings: 31.4 ± 6.0%, n = 22 fish; t-test, p = 0.023). Transgenics expressing a dominant negative construct (elavl3:shank3b^ΔC^) also spent more time next to the fish movie (elavl3:shank3b^ΔC^: 75.1 ± 2.8%, n = 10 fish; Wt siblings: 43.6 ± 8.9%, n = 14; t-test, p = 0.008).

We next analyzed the behavioral and neuronal activity of shank3b^−/−^ fish in the VR during normal visuomotor coupling. The fraction of time engaged in swimming behavior was slightly but not significantly lower in shank3b^−/−^ fish (23.5 ± 2.9 %, n = 11) as compared to wt siblings (27.6 ± 4.0 %, n = 10; t-test, p = 0.41; Figure S4C), consistent with the observations during free swimming. The mean AP probability, however, was significantly decreased in rDc and Dl of shank3b^−/−^ fish (746 and 2105 neurons in rDc and Dl, respectively) as compared to wt siblings (912 and 2326 neurons, respectively; rDc: p = 2.3e-34; Dl: p = 1.9e-32; Wilcoxon rank sum test; Figure 6A). The reduction was also significant when the mean neuronal activity of individual animals was compared (Figure 6B; shank3b^−/−^: n= 11; wt sibling: n = 10; t-test, rDc: p = 0.024; Dl: p = 0.049). In addition, we observed an increase in the fraction of swim-modulated neurons in Dm, cDc and Dl but not in rDc (Figure S4D). Overall, adult zebrafish with homozygous shank3b mutations therefore showed reduced neuronal activity in rDc and Dl during spontaneous behavior despite similar swimming activity.

**Figure 6:**
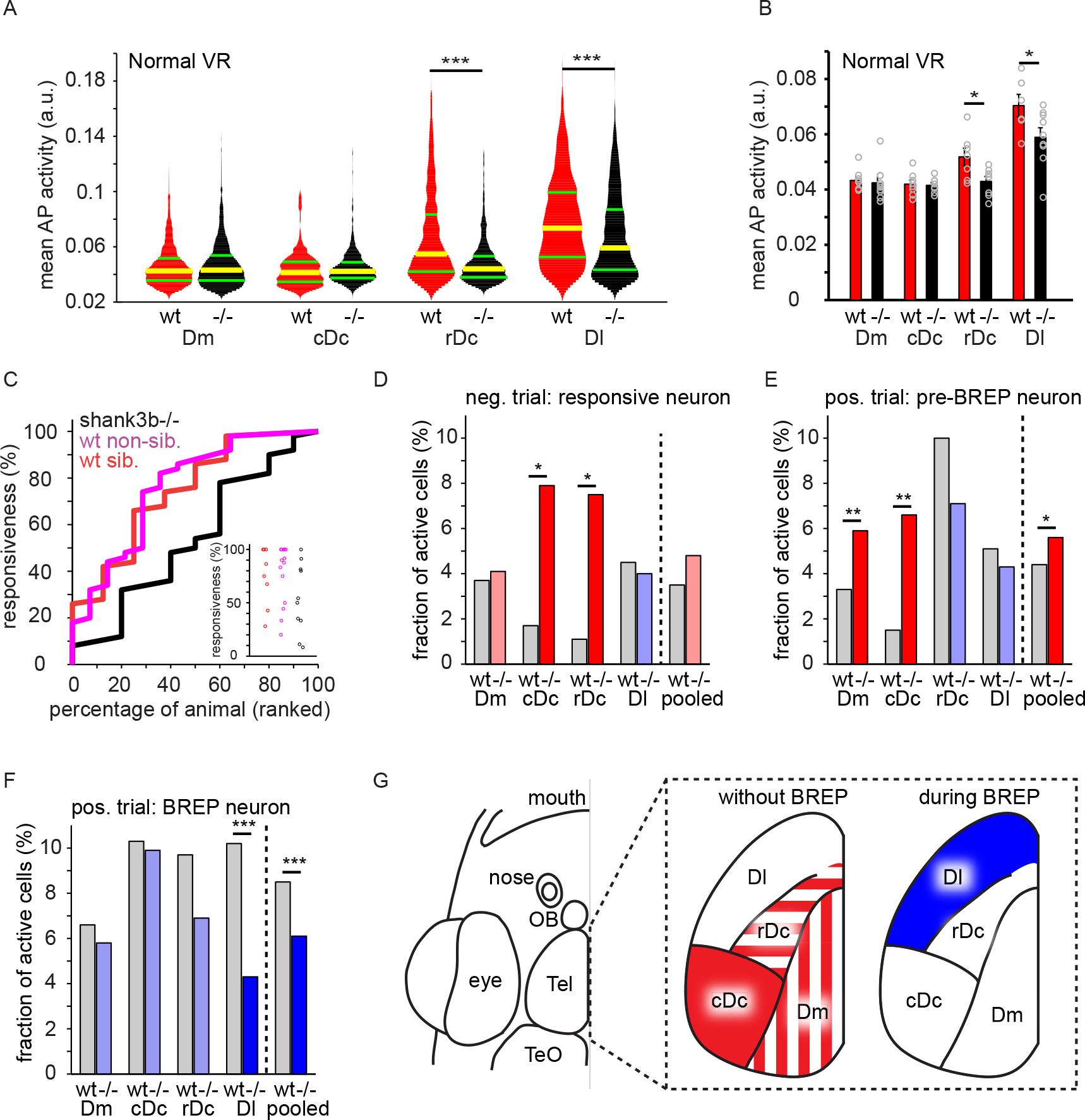
Perturbation of Visuomotor Mismatch Signals by Shank3b Mutation. (A) Shank3b^−/−^ fish showed reduced neuronal activity in rDc and Dl during spontaneous behavior in a VR with normal visuomotor coupling (p = 2.3e-34 and p = 1.9e-32, respectively, Wilcoxon rank sum test). Pooled results from 11 shank3b^−/−^ fish (Dm, 2275 cells, cDc, 832 cells; rDc, 746 cells; Dl, 2105 cells) and 10 wt siblings (Dm, 2411 cells; cDc, 498 cells; rDc, 912 cells; Dl 2326 cells). Yellow and green bars indicate median and interquartile range, respectively. (B) Mean AP probability in different brain regions and animals (shank3b^−/−^: n = 11; wt: n = 10). In shank3b^−/−^ fish, mean activity was significantly lower in rDc and Dl (p = 0.024 and p = 0.049, respectively; t-test). (C) Cumulative distribution of behavioral responsiveness to VR perturbations in shank3b^−/−^ fish (n = 10), wt siblings (n = 8), and wt non-siblings (n = 14). Responsiveness was defined as the fraction of positive trials in each animal. Inset shows responsiveness in each individual. (D) Fraction of neurons responding after the onset of a VR perturbation during negative trials as a function of brain area and genotype. In shank3b^−/−^ fish, the fraction of responsive neurons was significantly higher than in wt fish in telencephalic areas cDc and rDc. Wt siblings: Dm, 3.7% of 669 cells; cDc, 1.7% of 121 cells; rDc, 1.1% of 175 cells; Dl, 4.5% of 426 cells. Shank3b^−/−^: Dm, 4.1% of 459 cells; cDc, 7.9% of 202 cells; rDc, 7.5% of 133 cells; Dl, 4.0% of 754 cells. Red (blue) color depicts higher (lower) values in shank3b^−/−^ fish as compared to wt siblings. Strong colors depict statistically significant differences. (E) Fraction of neurons responding during pre-BREP period of positive trials as a function of brain area and genotype (neurons responding after perturbation onset but before onset of the BREP). In shank3b^−/−^ fish, the fraction of responsive neurons was significantly higher than in wt fish in telencephalic areas Dm and cDc. The fraction of responsive neurons was also significantly higher when neurons were pooled across all brain areas (right). Wt siblings: Dm, 6.6% of 1357 cells; cDc, 10.3% of 271 cells; rDc, 9.7% of 259 cells; Dl, 10.2% of 1101 cells. Shank3b^−/−^: Dm, 5.8% of 625 cells; cDc, 9.9% of 467 cells; rDc, 6.9% of 437 cells; Dl, 4.3% of 1035 cells. Colors as in D. (F) Fraction of neurons that started to respond during the BREP period of positive trials as a function of brain area and genotype. In shank3b^−/−^ fish, the fraction of responsive neurons was significantly lower in Dl. The fraction of responsive neurons was also significantly lower when neurons were pooled across all brain areas (right). Wt siblings: Dm, 3.3% of 1357 cells; cDc, 1.5% of 271 cells; rDc, 10.0% of 259 cells; Dl, 5.1% of 1101 cells. Shank3b^−/−^ mutants: Dm, 5.9% of 625 cells; cDc, 6.6% of 467 cells; rDc, 7.1% of 437 cells; Dl, 4.3% of 1035 cells. Colors as in D. (G) Summary: differences in responses to VR perturbations between shank3b^−/−^ fish and wt siblings. Left: during the pre-BREP period or in negative trials (no behavioral change); right: during the BREP. During the pre-BREP period, activity in shank3b^−/−^ fish was significantly increased in cDc (in positive and negative trials; solid color), in rDc (in negative trials; striped), and in Dm (in positive trials; striped). During the BREP period, activity in shank3b^−/−^ fish was significantly lower in Dl.

We next investigated how the behavior and neuronal activity of shank3b^−/−^ fish were affected by a left-right reversal of visuomotor coupling. Shank3b^−/−^ fish exhibited a BREP in 61% of the perturbation trials (n = 200 trials), significantly less than wt siblings (71%, n = 141 trials) and wt non-siblings (75%, n = 113 trials; Figure S4E, X^2^ test between shank3b−/− and pooled wt, p = 0.015 after Bonfferoni correction). Consistent with this observation, the fraction of highly responsive animals (fish exhibiting a BREP in >70% of trails) was lower in shank3b^−/−^ fish (4 out of 10; 40%) than in wt siblings (5 out of 8 fish; 62.5%) and wt non-siblings (10 out of 14 fish; 71.4%; Figure 6C) although this effect did not reach statistical significance (t-test, p = 0.06).

We next compared neural activity in response to visuomotor perturbations between shank3b^−/−^ and wt fish. Because the motor output during the positive and negative behavioral trials were qualitatively different we analyzed the neuronal responses separately under each behavioral condition. During negative trials, we found very few responsive neurons throughout the dorsal forebrain in wt siblings (Dm 3.7% of 669 cells; cDc 1.7% of 121 cells; rDc, 1.1% of 175 cells; and Dl 4.5% of 426 cells, Figure 6D). In shank3b^−/−^ fish, in contrast, a substantial fraction of neurons were responsive in the Dc regions (cDc: 7.9% of 202 cells, p=0.033; rDc: 7.5% of 133 cells, p=0.010, X^2^ test).

During positive trials, we further distinguished between BREP and pre-BREP neurons. The fraction of BREP neurons was lower in shank3b^−/−^ fish than in wt siblings in all brain areas, particularly in Dl (Figure 6F, shank3b^−/−^: 4.3% of 1035 cells; wt siblings: 10.2% of 1001 cells; X^2^ test, p = 3.9e-7). As a result, the overall fraction of BREP neurons in dorsal forebrain was significantly decreased (6.1% of 2564 cells in shank3b^−/−^ vs. 8.5% of 2988 cells in wt siblings, X^2^ test, p = 8.9e-4). The fraction of pre-BREP neurons, in contrast, was increased in shank3b^−/−^ fish (Figure 6E, 5.6% of 2564 cells vs. 4.4% of 2988 cells in wt siblings, p = 0.047, X^2^ test). This overall increase in pre-BREP activity was due to significant increases in the fraction of pre-BREP neurons in Dm and cDc (Dm: 5.9% of 625 cells in mutant fish and 3.3% of 1357 cells in wt siblings, p = 0.0098, X^2^ test. cDc: 6.6% of 467 cells in mutant fish and 1.5% of 271 cells in wt siblings, p = 0.0027, X^2^ test). In area Dm, we further observed that the activity pattern of pre-BREP neurons was more heterogeneous than in wt siblings (Figure S4F).

In summary, we found that responses to sensorimotor mismatch before the BREP were increased in Dm and in parts of Dc of shank3b^−/−^ fish as compared to wt siblings. In negative trials, shank3b^−/−^ fish showed a similar pattern of increased responses to sensorimotor mismatch without an apparent behavioral response. The activity of neurons responding during the BREP, in contrast, was lower in shank3b^−/−^ fish than in wt siblings, particularly in Dl. Hence, responses to sensorimotor mismatch exhibited distinct differences between shank3b^−/−^ and wt fish that appeared to be complementary during the pre-BREP and BREP phases (Figure 6G). The shank3b^−/−^ mutation therefore had multiple effects on prediction error signaling across different brain areas.

## Discussion

We developed techniques to measure neuronal activity in adult zebrafish during behavior in a VR, manipulated sensory feedback during motor behavior, and found prominent responses to sensorimotor mismatch in the zebrafish telencephalon. These results show that prediction errors are encoded by distributed populations of telencephalic neurons, consistent with the assumption that predictive processing is a widespread computation. Spontaneous activity and prediction error signals were altered in zebrafish carrying a mutation in shank3b, a gene whose mammalian ortholog has been linked to Phelan-McDermid syndrome, a form of autism (Phelan and McDermid, 2011). Prominent differences between wt and mutant fish included an enhanced response to sensorimotor mismatch in telencephalic areas Dm and Dc, which are thought to be homologous to mammalian basolateral amygdala and isocortex, respectively. These results are consistent with the hypothesis that conditions related to autism include an imbalance between sensory input and internal predictions.

### Neuronal Responses for Predictive Processing in the Telencephalon of Zebrafish

We developed a head fixation procedure and a high-resolution, closed loop VR that allowed us to perform multiphoton calcium imaging in adult zebrafish through the intact skull during visuomotor behaviors. This approach provides new opportunities to study the neural basis of behaviors that are rudimentary or absent in zebrafish larvae, including associative learning and social behavior. VR-based approaches are particularly powerful to analyze neural activity related to predictive processing because they allow the experimenter to decouple visual input from motor output. Using this approach we found that brief inversions of sensorimotor coupling evoked strong behavioral responses (“BREPs”) that consisted of unilateral tail bends followed by vigorous irregular tail movements. The initial unilateral maneuvers resembled orienting movements known as J-turns (McElligott & O’Malley, 2005; Bianco et al., 2011), suggesting that fish initially tried to correct their swimming course in response to unexpected visual input. Later phases of the BREP may then represent struggling behavior following a failure in course control.

The BREP was preceded by a “pre-BREP” period that followed the visuomotor perturbation but preceded the transition in motor behavior. During this period, fish still performed normal swim bouts although sensorimotor coupling was already reversed. Fish therefore experienced a mismatch between the expected and the actual sensory feedback, which most likely triggered the subsequent behavioral response. The level of neural activity during both the pre-BREP and the BREP period clearly exceeded the activity associated with strong spontaneous swim events, indicating that most of the activity was evoked by the sensorimotor perturbation.

Multiple observations indicate that pre-BREP neurons encode the mismatch between the predicted and the experienced sensory input. First, tail movements during the pre-BREP period were not significantly different from the period before the perturbation, indicating that the pre-BREP response is not related to a change in motor output. Second, the activity level of pre-BREP neurons predicted the occurrence of a subsequent BREP, consistent with the hypothesis that pre-BREP neurons encode an error signal that triggers error correction behavior. Third, the activity of pre-BREP neurons was not enhanced before spontaneous swim events of long duration, indicating that it does not represent preparatory motor activity. This conclusion is further supported by the observation that the activity of pre-BREP neurons was elevated during, rather than before, individual swim bouts. Fourth, the activity of many pre-BREP neurons was coupled to swim events during spontaneous motor behavior, consistent with the assumption that these neurons respond to sensory feedback during motor output. Fifth, while motor output during the pre-BREP period was indistinguishable from motor output during spontaneous swims, swim-coupled activity was greatly enhanced when visual feedback was perturbed. This observation indicates directly that pre-BREP responses do not reflect motor output but the mismatch between the expected and the experienced sensory feedback. We therefore conclude that neuronal responses during the pre-BREP period most likely reflect prediction error (“mismatch”) signals.

Responses of BREP neurons also depended on sensorimotor mismatch but occurred during enhanced motor output. One possibility is therefore that BREP activity is involved in motor control. However, the activity of BREP neurons was much higher during BREPs than during spontaneous swimming of similar intensity without VR perturbation, suggesting that at least some BREP activity was not motor-related. Another possibility is that BREP activity reflects a change in emotional state that accompanies the behavioral transition. Conceivably, the failure to control the course of swimming may result in anxiety or frustration. A third possibility is that BREP activity represents additional error signals. Such error signals are expected to occur during the BREP because the mismatch between the vigorous motor output and the resulting sensory input is very large. Responses of BREP neurons are therefore likely to reflect multiple processes including the signaling of sensorimotor prediction errors.

Experiments using a VR in mice revealed that a large fraction of neurons in visual cortex respond to perturbations of sensorimotor coupling. Responses of these “mismatch” neurons are very similar to pre-BREP, and possibly BREP, responses in the telencephalon of zebrafish. Hence, mismatch responses representing sensorimotor error signals are prominent not only in mammals but also in teleosts. We further found that mismatch responses are distributed across multiple brain areas. These responses may be generated locally in each brain area by comparisons between sensory input and internal predictions, or they may be computed in one area and transmitted to others by projection neurons. Independent of the mechanism that generates mismatch responses, their widespread occurrence demonstrates that information about sensorimotor prediction errors is available throughout large parts of the telencephalon. These results are consistent with the hypothesis that predictive processing is an important computational principle in the brain (Keller and Mrsic Flogel 2018).

### Aberrant Processing of Prediction Errors in shank3b^−/−^ fish

An imbalance between sensory input and an internal prediction has been proposed to create a sensory overload that underlies key symptoms of autism such as repetitive behaviors (Pellicano and Burr, 2012; Friston et al., 2013; Lawson et al., 2014; Lawson et al., 2017; Sinha et al., 2014). This hypothesis directly predicts that mutations linked to autism spectrum disorders result in abnormal prediction error signals. In sensory brain areas of autism models it may be expected that prediction error signals are enhanced because excitatory sensory input is not cancelled by an inhibitory prediction of appropriate specificity and strength. However, considering network interactions between brain areas, an imbalance between sensory input and prediction may also result in more complex and region-specific changes in neuronal activity. To explore this question we examined fish with a mutation in shank3b^−/−^. These fish showed an enhanced attraction to movies of conspecifics that was phenocopied by the overexpression of the dominant-negative shank3b isoform shank3b^ΔC^, indicating that specific effects on behavior were caused by the disruption of shank3b function. Consistent with this conclusion, a previous study reported that a mutation in shank3b resulted in reduced social interactions and enhanced repetitive behaviors in adult zebrafish (Liu et al, 2018).

Our approach allowed us to perform detailed comparisons between activity patterns in shank3b^−/−^ fish and wt siblings. Neuronal responses to sensorimotor mismatch in shank3b^−/−^ fish were enhanced before the BREP but reduced after BREP onset in complementary brain areas (Figure 6G). While responses during the BREP may be associated with different processes, pre-BREP responses are likely to represent prediction errors. The enhanced pre-BREP activity in shank3b^−/−^ fish is therefore consistent with the hypothesis that a mutation in shank3b increases prediction error signals due to an imbalance between sensory input and internal predictions. Enhanced pre-BREP activity was observed specifically in brain areas related to isocortex (Dc) and to the basolateral amygdala (Dm). These results are consistent with the hypothesis that mutations related to autism modify predictive processing and provide a starting point to explore how aberrant predictive processing affects behavior.

## Acknowledgements

We thank Georg Keller for comments on the manuscript; Paul Argast, Peter Buchmann and Pawel Zmarz for technical support, Andrew Prendergast and Claire Wyart for sharing NeuroD:GCaMP6f fish, Jessica Garver, Stacey Gearin and Stephanie Wiessner for excellent technical assistance; Otto Fajardo for support during the initial phase of the project, and the Friedrich group for stimulating discussions. This work was supported by Novartis Research Foundation (R.W.F.), by a Novartis Institutes for Biomedical Research Presidential Postdoctoral Fellowship (K.H.), by the European Research Council (ERC) under the European Union’s Horizon 2020 research and innovation programme (grant agreement No 742576), and by a fellowship from the Boehringer Ingelheim Fonds (P.R.).

## Author Contributions

K.H. developed the methodology, designed and performed experiments, analyzed data, and wrote the manuscript. P.R. developed the methodology and wrote the manuscript. M.S. and F.S. provided reagents including transgenic fish. K.K. performed experiments using freely behaving fish. T.B. supervised the project. R.W.F. supervised the project, analyzed data and wrote the manuscript.

## METHODS

### EXPERIMENTAL MODEL AND SUBJECT DETAILS

Experiments were performed in adult (5 – 21 months old) zebrafish (*Danio rerio*). Fish were raised and kept under standard laboratory conditions (26–27°C; 13 h/11 h light/dark cycle). Shank3b^−/−^ fish were obtained from the Zebrafish International Resource Center (ZIRC, allele sa2155). To generate Tg(elavl3:shank3bΔC) fish, the sequence of shank3b was truncated as described for the mouse shank3 gene (Durand et al, 2007), synthesized, cloned into an expression vector from the tol2kit containing the elavl3 promoter (http://tol2kit.genetics.utah.edu), and introduced into embryos using standard procedures. Assays using freely swimming fish were performed with 16 shank3b^−/−^ fish and 22 wt siblings, and with 10 dn(elavl3:shank3bΔC) fish and 14 wt siblings. Genders were equally represented. For calcium imaging experiments on head-fixed animals, a stable transgenic line expressing the fluorescent calcium sensor GCaMP6f under the control of a fragment of the neuroD promotor (Tg(NeuroD:GcaMP6f)icm05, Rupprecht et al., 2016) in a nacre background was used. These fish had no iridophores over the forebrain, improving optical access through the intact skull. Care was taken not to overfeed fish in order to reduce accumulation of lipid droplets between the skull and the brain. In total, the behavior of 37 head-fixed animals was examined in the closed-loop VR. Simultaneous calcium imaging was performed in 32 of these animals which included 10 shank3b^−/−^ fish, 8 wt siblings and 14 wt non-siblings (11 males and 21 females). All mutant and transgenic animals were generated by incrossing heterozygous mutants and transgenics, respectively. The offspring was then genotyped to identify homozygous mutants or transgenics and wt siblings for experiments. All experiments were approved by the Veterinary Department of the Canton Basel-Stadt (Switzerland).

## METHOD DETAILS

### Overview of the experimental procedure

Under anesthesia, light-weight head bars were attached to the skull of the fish using tissue glue and dental cement. Head bars were then glued to mounting posts and fish were allowed to recover from anesthesia. Fish were transferred to the VR environment underneath the two-photon microscope and habituated to the VR for 20-30 min in closed loop. During this period, the linear and angular feedback gain were adjusted manually until the animal exhibited quasi-periodic swimming behavior. Calcium imaging was performed in sessions that lasted 30 min. The field of view in each session covered one or multiple forebrain regions. VR perturbations (left-right reversal) were introduced either automatically every 7 min, each lasting for 10 s, or manually every 2 - 7 min, each lasting 5 - 10 s. In each fish, 2 - 8 calcium imaging sessions were performed that were separated by 15 min. During the time between sessions, calcium imaging data were converted to tiff image series and a new location for neural recording was identified while animals continued to behave in a closed-loop VR.

### Head-fixation of adult zebrafish for in vivo imaging

Fish were anaesthetized in 0.03% tricaine methanesulfonate (MS-222), wrapped in moist tissue and placed under a dissection microscope. To maintain anesthesia during surgery, 0.01% MS222 was continuously delivered into the mouth through a small cannula. The surgical procedure took approximately 50 min including gluing head bars to the skull (30 min), removing the skin above the forebrain to improve optical access (5 min), and gluing head bars to the mounting posts (15 min). After successful surgery basic behaviors of the animal including saccades, locomotion, feeding and social affiliation are not impaired (Movie S3). Detailed steps of the surgery are described below:

1. Anaesthetize the animal using 0.03 % MS-222 in fish tank water.
2. Wrap the anesthetized fish in a piece of moist paper tissue (e.g., Kimwipe) starting from the tip of the tail to ca. 1 mm caudal to the gills. Wrapping should be tight enough to prevent sliding of the animal out of the tissue. The tissue should not be too close to the gills to prevent the accumulation of water flowing over the gills. Wrap an appropriately sized plastic tubing with a longitudinal slit (e.g. a medium-sized heat-shrink tubing cut open) around the tissue to cover the fish from the end of the tail to ca. 2 mm caudal to the gills. This sleeve provides mechanical support to the fish, ensuring that the body remains straight throughout the surgery even when the head extends into the air (Figure S1D).
3. The attachment site of the head bars (“gluing site”) is a triangular area lateral to the cerebellum and above the gills (ca. 0.5 mm^2^). The skull is directly underneath the skin at this location. Remove the skin covering the gluing sites by scratching against the skull with a scalpel and dry the skull.
4. Apply a small amount of tissue glue (3M Vetbond) at the center of the gluing site using a 10 μL pipette tip. The tissue glue should expand on the exposed skull and quickly cover the entire gluing site.
5. The L-shaped head bar (4 by 1.5 mm, 2.8 mg) is made from a syringe needle (BD Microlance, OD = 300 μm) and can be reused for multiple experiments. Insert the long arm of the head bar to a piece of clay, which is solid enough to hold the head bar before gluing but soft enough to detach from the head bar once it is glued to the skull. Attach the clay to a 3-axis micromanipulator for precise alignment of the head bar to the gluing site. The short arm of the head bar should be parallel to the midline of the head and the long arm should be horizontal.
6. Mix the powder (0.05 g) and liquid (400 μL) components of dental cement (Paladur, Kulzer), and apply the mixture to cover the gluing site including the head bar using a 20 μL pipette tip. Dental cement should not contact the gills, eyes or water throughout the surgery. Curing of the dental cement takes ca. 3 min. Glue the second head bar on the other side of the head. The long arms of the two head bars should roughly form a line perpendicular to the body axis.
7. Apply a small amount of dental cement to connect the two head bars. This “cross linking” blocks optical access to the midbrain but provides enhanced stability.
8. Remove the skin over the telencephalon using a scalpel and fine scissors to improve optical access for calcium imaging. Skin removal should be avoided at two sites where blood vessels are located (Figure S1E).
9. Glue long arms of head bars to a pair of solid posts using dental cement (0.15 g powder mixed with 640 μL liquid component). Full curing of the cement takes ca. 10 − 15 min.
10. Immerse the animal in fresh water. The fish should recover from anesthesia within 1 − 2 min.

### Tail tracking and inference of virtual movement

The tail of the head-fixed fish was illuminated from the side using an infrared light (805 nm, Roithner Lasertechnik, RLT80810MG) and monitored from below using a camera (Point Grey, Dragonfly2). Image acquisition (50 Hz) was controlled by a custom program written in LabVIEW (National Instruments). A short pass filter (<875 nm, Edmund Optics, #86-106) and a long pass filter (>700 nm, Edmund Optics, #43-949) were positioned in front of the camera to block infrared light from the femtosecond laser and visible light from the projectors, respectively. Tail movements were recorded at a resolution of 90 by 70 pixels (lateral x rostro-caudal). The backbone of the tail was analyzed in real-time (50 Hz) using a custom script written in MATLAB (The Mathworks) (modified from Huang et al., 2013).

The curvature and the undulation frequency of the caudal 1/3 of the tail was quantified to infer the swimming trajectory and update the VR. Tail curvature only affected the VR when the tail was in motion, which was determined by quantifying the mean pixel-wise difference between adjacent frames. A curved but inactive tail did therefore not result in an update of the VR.

Forward motion and turning were determined based on the zero-crossings and the asymmetry of tail curvature, respectively (Figure 1G). The speed of forward swims was thus determined by the undulation frequency of the caudal tail. Every zero-crossing of the tail curvature triggered a forward movement that decayed exponentially (exponential decay constant, τ = 190 ms). Thus, fast symmetric tail undulations were most efficient in driving forward swims in the VR.

The amplitude of turns was derived from the bending direction and the curvature of the caudal tail. In each video frame of the behavior recording, a right bend triggered a right turn in proportion to the tail curvature, followed by an exponential decay (τ = 190 ms). A sequence of unilateral tail bends was therefore most efficient in driving a large rotation in the VR. In contrast, fast symmetric tail undulations, i.e. forward swims, caused minimal angular updates of the VR.

The gain of the forward swims and turns in the VR was manually adjusted in each animal during the first 20-30 min of the experiment, prior to activity measurements. The goal of the gain adjustment was to obtain realistic swimming behavior without apparent signs of stress. After successful adjustment, animals did not struggle and performed controlled, periodic swims. Under these conditions, fish remained active and constantly performed calm periodic swims for more than 6 h.

### Virtual reality

A 3D model of the virtual reality was built using the modelling software Wings 3D. The virtual environment consisted of a cylindrical arena with two tunnels on opposite sides. The Python-based game engine Panda3D was used to update the location and angle of the virtual cameras in the VR. A cluster of three virtual cameras were used and each virtual camera captured a 60 ° by 60 ° (height by width) field of view side by side. Therefore the camera cluster captured a 60 ° by 180 ° (height by width) field of view, which was projected to the semi-hexagonal VR chamber. Forward swims translated into a forward movement of the virtual cameras and right (left) turns of the fish translated into clockwise (counter-clockwise) rotation of the virtual cameras. The VR also permitted the presentation of dynamic textures including movies at specific locations of the virtual environment but this feature was not systematically used in the present experiments. However, in some experiments, movies of moving objects were presented inside the tunnels.

Collision detection was included in the Panda3D code to keep the animal at a distance from visible boundaries to prevent pixilation of textures. Specifically, the virtual cameras capturing the field of view in the VR were enclosed by an invisible collision sphere, which would be pushed back by the invisible collision wall in the 3D model (Figure 1F). The camera cluster therefore stopped moving when it hit the wall head-on, or it slid along the wall smoothly when it struck it at an angle. Since the VR chamber was filled with water, the VR could not be projected onto the interior of a toroid screen (Harvey et al., 2009). Instead, the VR was projected onto back projection film (DILAND SCREEN, DGP) from the outside. The VR chamber was covered by this film and had a semi-hexagonal shape (10 cm per side, 8 cm high, made of polymethyl methacrylate). Three virtual cameras were introduced in the VR, each with a field of view of 40° by 20° (width x height). Together, the three cameras therefore projected a continuous VR texture that covered 120° in the horizontal direction. The VR texture captured by the three virtual cameras was projected by three projectors (AAXA P3, with RGB LEDs and LCoS display) onto the three sides of the semi-hexagonal tank. As described below, the red, green and blue LEDs of the projectors were gated by the line trigger signal (TTL) from the resonant scanner (Cambridge Technology, CRS 8 kHz) using a custom-built circuit (Leinweber et al., 2014) to achieve fast rise and fall times of the LED output.

### VR combined with two-photon microscopy

Adult zebrafish were head-fixed and positioned at the rear center of the water-filled semi-hexagonal chamber (water depth 7 cm), ca. 3 cm below the water surface. A 16x water-immersion objective with a working distance of 3 mm (Nikon, CF175 LWD 16xW) was used for two-photon imaging. A custom-designed water-proof sleeve allowed for immersion of the objective >3 cm below the water surface (Figure 1D). A custom-built multiphoton microscope with resonant scanners was used to acquire series of images with 512 x 512 pixels at a rate of 30 Hz using custom-written software based on Scanimage (Pologruto et al., 2003; Rupprecht et al., 2016). Because the photomultiplier was gated on alternating lines (see below), the effective resolution was 512 x 256 pixels at 30 Hz.

GCaMP6f fluorescence was imaged through the intact skull using an excitation wavelength of 920 nm with a power of 50 mW at the sample. In contrast to a head-fixed rodent, where the front lens of the objective lens is usually surrounded by the skull and the metal-made head plate (Dombeck et al., 2007), the objective lens for imaging of head-fixed zebrafish was entirely outside of the fish head. As a consequence, a substantial amount of photons from the virtual reality entered the objective lens. In order to avoid contamination of fluorescence measurements by these photons we temporally separated calcium imaging from the VR. This was achieved by gating the light-emitting diodes of the projector and the fluorescence-detecting gated photon multiplier tube (H11706P-40, Hamamatsu) in a non-overlapping manner. The TTL line clock of the resonant scanner (Cambridge Technology, CRS 8 kHz) was used as a gating signal to switch between VR illumination and fluorescence detection on alternating lines (Figure 1D). This line-by-line triggering of the electronics at high speed required a data acquisition board with a retriggering function (National Instruments, PCIe-6321). The projector LEDs were turned ON for 24 μs during the turnaround period of the resonant scanning. The shutter circuit was switched to a low gain during this period. To further reduce the contamination of calcium imaging by photon noise from the VR, a notch filter (514.5/25 nm, OD 4, Edmund Optics) was positioned in front of each projector and an additional band-pass filter (510/50 nm, Chroma) in front of the PMT. In-between imaging sessions, the resonant scanner was not scanning and the LEDs were triggered by another 8 kHz TTL signal to maintain a constant virtual environment for the fish. To detect the precise temporal frame-to-frame correspondence between behavioral and neuronal activity, the frame count of the two-photon recording (30 Hz) was stored in each video frame of the behavior recording (50 fps).

### Behavioral assay for freely-swimming adult zebrafish

The behavior of individual free-swimming fish was recorded in a rectangular tank (L*W*H = 20*10*15 cm, water depth 10 cm) for one hour at 10 frames per second (Figure 5A). The long walls of the tank were made of white PVC with matt surface. The short walls were made of anti-reflection glass (LUXAR, glastroesch) to prevent interactions of fish with their mirror image. The bottom of the tank was covered with a diffusor to prevent reflection. An infrared LED panel was used to illuminate the tank from below for video imaging from above. Swim velocity (Figure S4A & S4B) was calculated by measuring the displacement of the center of mass of the body during 2 s intervals. To test the animal’s response to controlled visual access of conspecifics, a movie of two females and one male was presented to the animal for 20 s using a LCD monitor that was placed immediately behind one of the short walls.

## QUANTIFICATION AND STATISTICAL ANALYSIS

### Definition of brain regions in the dorsal pallium

Anatomical definitions of canonical subdivisions of the dorsal pallium in adult zebrafish vary somewhat between previous studies (Wullimann et al., 1996; Mueller et al., 2011; Aoki et al, 2013; Ganz et al., 2014). We adopted the definitions of Aoki et al. (2013) with minor modifications because this definition best matched landmarks in the fluorescence images of the dorsal telencephalon from NeuroD:GCaMP6f transgenic fish. At the end of the experiment, two-photon image stacks of the dorsal pallium were acquired in each fish and forebrain regions were delineated manually based on anatomical features. Dm was distinctly separated from other forebrain regions by the sulcus ypsilonformis. cDc was separated from rDc by a boundary that was visible in the NeuroD:GCaMP6f expression (Figure S3). This boundary appears to correspond to a boundary in parvalbumin expression (Aoki et al, 2013). Dl was lateral to rDc at the rostral end of the forebrain but covered rDc at a more caudal position (Figure S3, also Mueller et al., 2011).

### Detection of the onset of swim events

To measure the onset of swim events, the mean of absolute difference in pixel values between adjacent video frames (50 Hz, 90 by 70 pixels per frame, covering the 2/3 caudal part of the tail) was calculated. This analysis yields a measure of the intensity of tail movements as a function of (Figure 2D, Figure 3C - F). The trace was binarized using a threshold of three standard deviations, calculated from the lower half of the intensity distribution. Binarized events shorter than 100 ms were removed and events closer than 100 ms were fused. The onset of swim events was then determined by threshold crossings (Figure 2D, black star signs).

The onset of BREPs was manually determined by inspection of the behavioral video. Fish typically responded to VR perturbations with increased motor output. It started with consecutive unilateral bends of the caudal tail followed by vigorous and often bilateral movements of the whole tail, including the rostral trunk of the body (Movie S2). In rare cases an abrupt transition in tail beat structure was not detectable. BREP onset was then set to the 3^rd^ tail flick after the VR perturbation.

### Post-processing of calcium imaging data and extraction of neuronal ROIs

For each session, a single optical plane was recorded for 30 min at 30 fps, resulting in 54000 frames per plane. The images were separated into 7 files with 8000 frames per file (267 seconds) except the last file which contained 6000 frames. For each file, a 2D Gaussian filter (standard deviation in x- and y-direction = 1 pixel) was applied, followed by a maximum projection in the time dimension. The resulting map highlighted the position of neurons that had been active during the recording period.

Two types of motion artifacts were observed. First, in rare cases, struggling behavior introduced a transient movement of the brain in both lateral (x, y) and axial (z) directions, lasting for 0-2 s. After the struggle, the brain sample typically returned to the original z-position which allowed for continued imaging of the same cells. These movement artifacts would result in “blurring” of the maximum projection image. Files containing such artifacts were therefore excluded from the analysis. The second motion artifact was a slow, lateral drift (<1 μm/min) towards the rostral direction. Because active neurons were sparsely distributed in the forebrain (except in some subregions of Dl), this slow drift did not interfere with the analysis of neuronal responses triggered by the VR perturbation, which lasted for ~10 seconds.

In most analyses, ROIs were drawn independently in different files and treated as different neurons. It is therefore possible that the same neuron appears more than once in the analysis when ROIs in different files represent the same neuron. In matrices representing the activity of neurons as a function of time, we estimate that each neuron was represented, on average, by 2.9 entries. For the analysis of the same pre-BREP neurons during positive and negative trials (Figure 4G, 4H), however, activity traces were concatenated for neurons that were identified reliably across all seven files of a session. This ensured that the same neurons were analyzed during positive and negative trials.

ROIs for single neurons were selected manually in ImageJ (Schneider et al., 2012). Time traces were averaged across all pixels for each ROI. For calculation of ΔF/F_0_ traces, F_0_ was determined as the mean of the 25 % lowest percentile of the fluorescence trace. To improve the temporal resolution of the neuronal activity traces, ΔF/F_0_ traces were deconvolved to extract relative action potential (AP) probabilities using the Elephant algorithm based on machine learning (Berens et al., 2018; https://git.io/vNbsz).

### Modulation of neuronal activity by swimming

To determine whether a neuron was positively modulated by swim events, the onsets of swim events were detected within the 267 s of each recording file. For each cell and for each swim event, the difference in AP probability Δp between 0.5 s time windows before and after swim onset was calculated, resulting in a distribution of swim-triggered Δp values for each cell. This distribution was compared to the distribution triggered on 500 random time points on the same activity trace using a one-sided t-test. Swim modulated neurons were defined by p < 0.05.

### Modulation of neuronal activity by VR perturbation

To determine whether a neuron was responsive to the VR perturbation, the difference in ΔF/F between 10 s time window before and after perturbation onset was calculated. The response value was compared to the response distribution triggered by 500 random time points on the same activity trace. The cell was considered responsive in this perturbation trial when the response value exceeded the mean by two standard deviations of the randomly sampled distribution.

To identify responsive neurons during spontaneous strong swims (Figure 4B, black), a time interval was randomly sampled from the pre-BREP period and paired to a spontaneous strong swim to define the “onset of perturbation”. This artificial onset was used to identify responsive neurons by the procedure described above.

The temporal response onset of an individual neuron (Figure 4B, red and green) was determined as follows. The activity trace from −10 s to +10s around the perturbation onset were smoothed using a median filter with a 150 ms time span. The lower 70 % of the activity trace before the perturbation were used to calculate the baseline and the standard deviation. Rising edges were detected from the post-perturbation activity trace using a threshold of 5 standard deviations. The first rising edge with a mean activity in the following 1 s higher than three standard deviations was defined as the first calcium transient. The combination of criteria for height (five standard deviations) and duration (three standard deviations for 1 s) reliably separated responses from noise. From the rising edge of the response event, a retrograde search was applied for the first time point when the activity was lower than 2 standard deviations. This time point was set as the response onset of the neuron to this respective perturbation.

## Supplemental information

**Figure S1.**
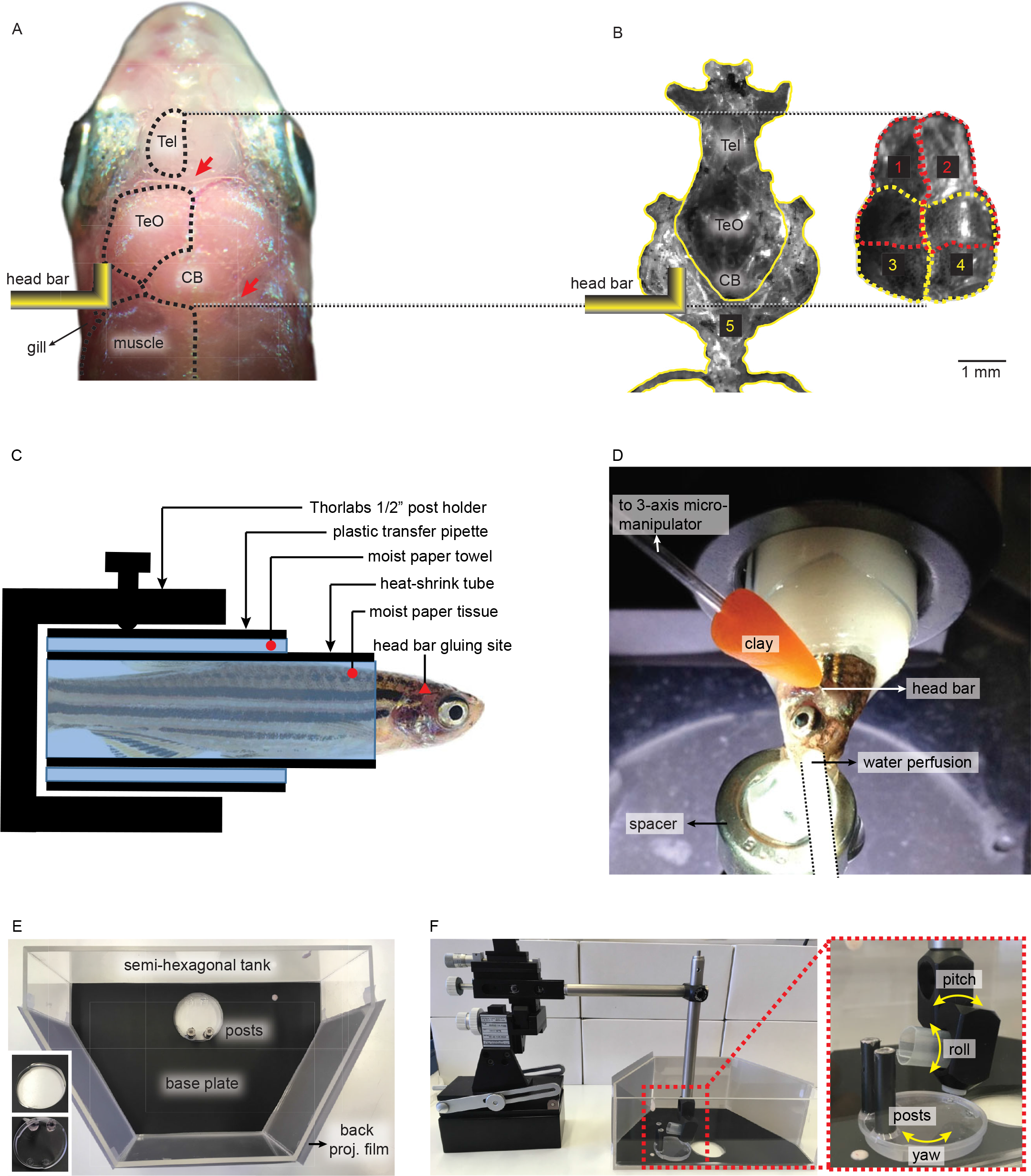
Head fixation of adult zebrafish. Related to Figure 1. A. Location for gluing L-shaped head bars. Skin removal should be avoided at two specific locations to prevent bleeding (red arrows). B. Five major sets of skull bones in adult zebrafish. Sets 1 - 4 are located on top of set 5. Tel: telencephalon, TeO: optic tectum, CB: cerebellum. C. Wrapping materials to keep skin moisturized and body stable during surgery. D. Setup to glue a head bar to the skull. E: Components of the VR chamber. Inset: mechanism to seal the mounting posts to the base plate. F. Tools to align the yaw, roll and pitch angles of the fish on the mounting posts.

**Figure S2.**
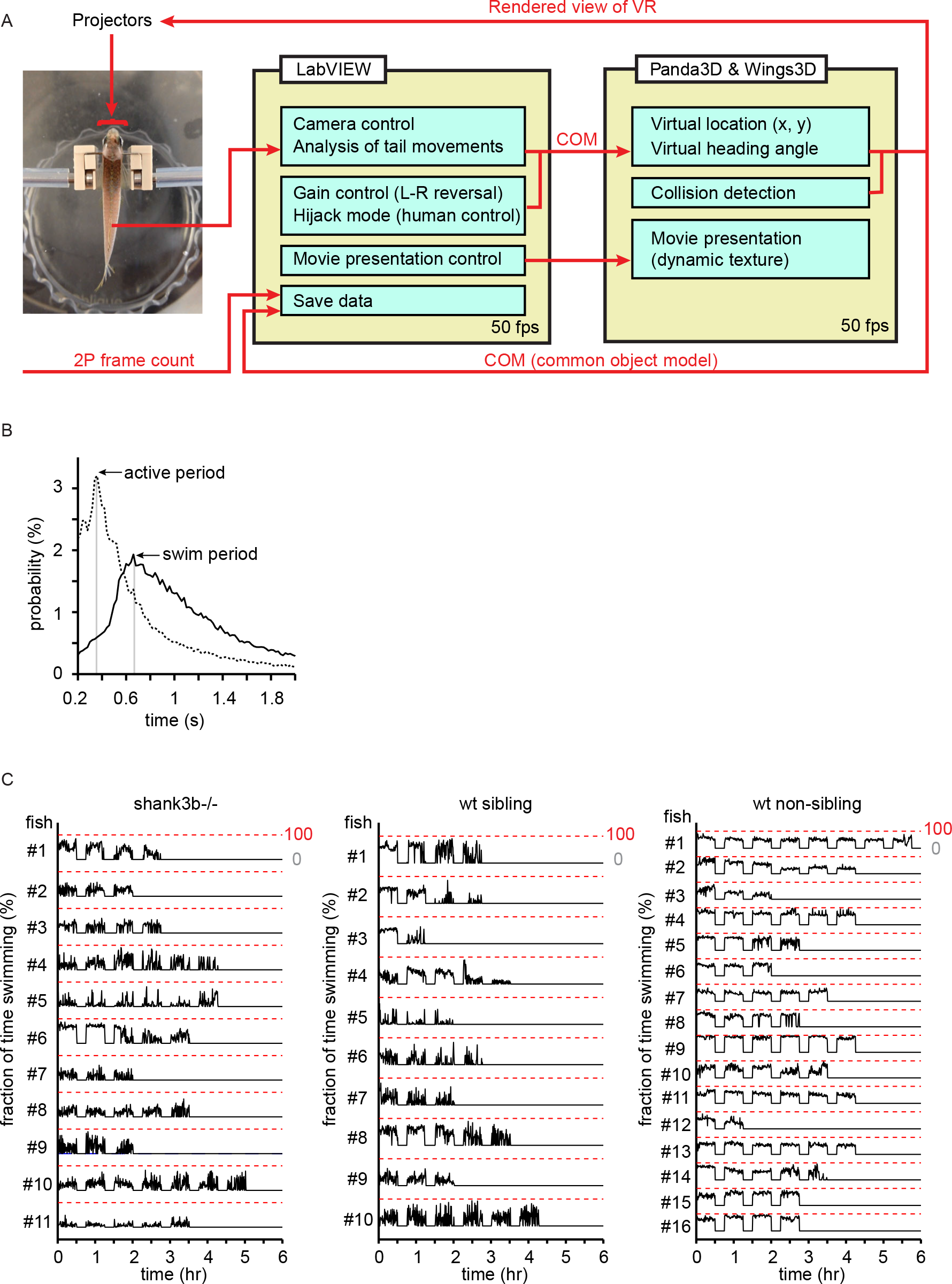
Swimming behavior of adult zebrafish in a closed-loop VR. Related to Figure 1. A. Workflow of a closed-loop VR. Photograph shows fish in a custom-made holder for reversible mounting. B. Temporal analysis of tail movements of head-fixed fish. Animals under head fixation exhibited quasi-periodical swim events (n = 37 fish, each recorded for at least 30 min). The duration of a swim event (swim period, mode = 0.66 s) consisted of a period of active tail movement (active period, mode = 0.36 s) followed by an inactive period. Tail movement was measured by the mean of absolute pixel-wise difference in adjacent frames of behavioral recordings. C. Head-fixed adult zebrafish remained behaviorally active for hours in a closed loop VR. Successive recording periods (30 min) were separated by non-recording periods (15 min) but a closed-loop VR was maintained throughout both periods. Swim probability was measured in 30 s time windows during recording periods. Stretches of zero motility correspond to non-recording periods.

**Figure S3.**
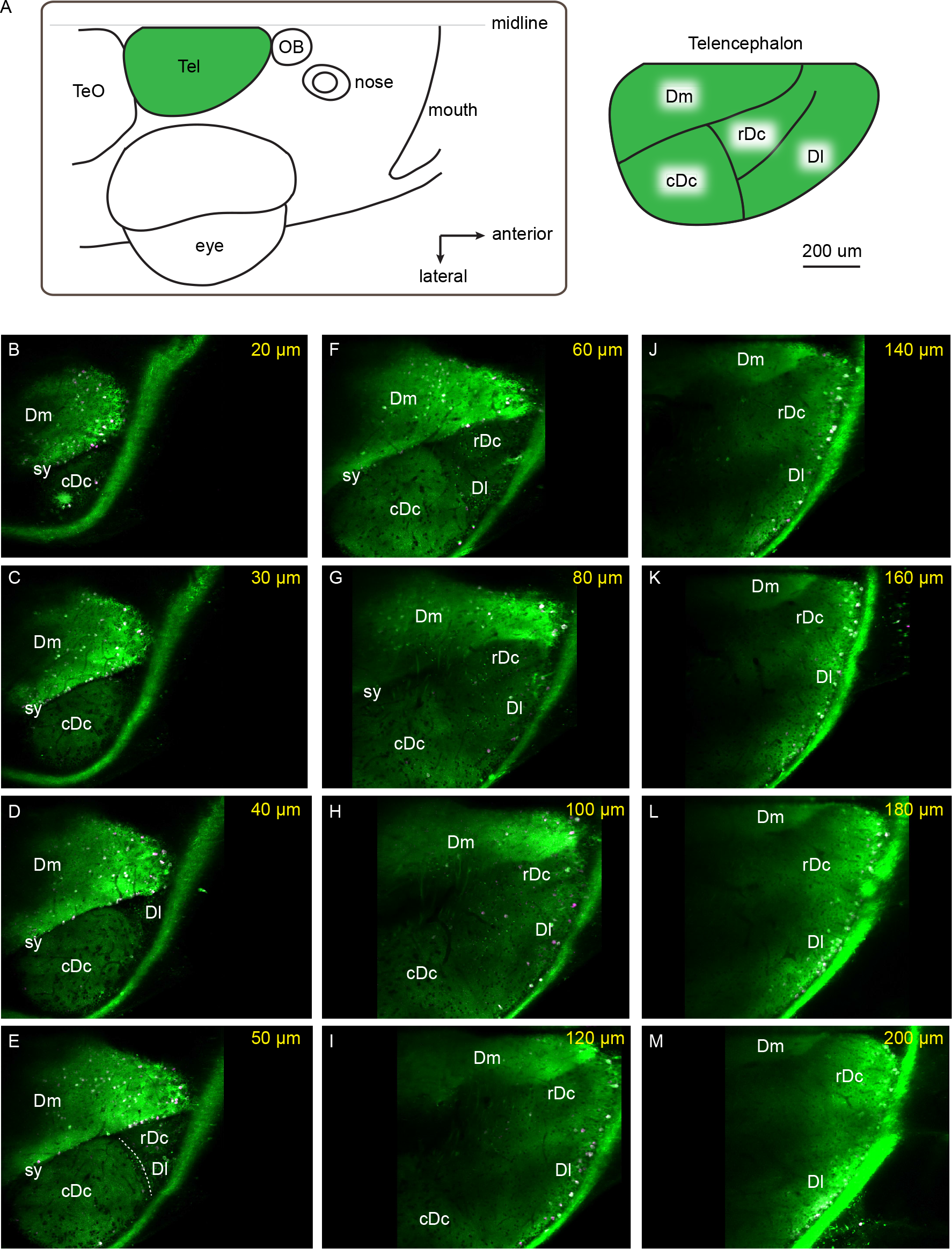
Expression of NeuroD:GCaMP6f in the dorsal telencephalon. Related to Figure 2. A. Schematic showing the location of the dorsal telencephalon (left) and its subdivisions (right). Tel: telencephalon; TeO: optic tectum; OB: olfactory bulb. B – M. NeuroD:GCaMP6f expression in a dorsal-to-ventral series of multiphoton images acquired through the intact skull. Green: raw fluorescence. Magenta: neuronal activity determined by maximum projection of pixel values in a time series of mean-subtracted images. Labels denote telencephalic brain areas. Dm is separated from Dc regions by the sulcus of ypsilonformis (sy). cDc was separated from rDc and Dl by an obvious anatomical boundary (dashed line in E). Dl is lateral to rDc at the rostral end of the forebrain but extends dorsally and covers rDc at caudal locations (D - F). z = 0 μm is defined by the dorsal surface of Dm.

**Figure S4.**
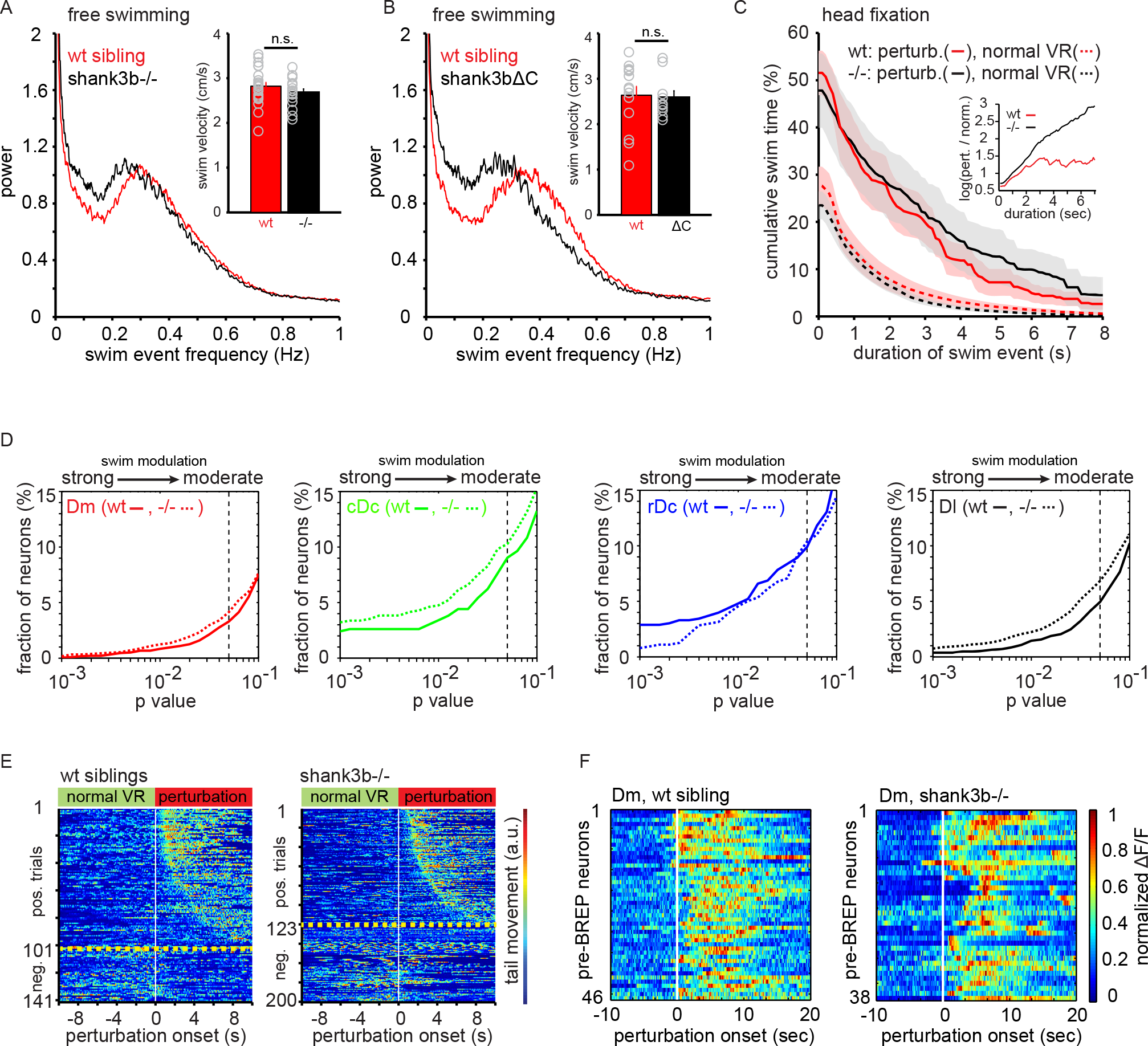
Behavioral and neural phenotypes of shank3b−/− fish. Related to Figures 5 and 6. A. Power spectral analysis of swim velocity traces from freely swimming fish. The mode of swim frequency in shank3b−/− fish (0.25 Hz) was slightly lower than in wt siblings (0.30 Hz). Inset: mean swim velocity was indistinguishable between mutants (2.67 ± 0.08 cm/s, mean ± SEM) and wt siblings (2.82 ± 0.09 cm/s). B. The mode of swim frequency in Dn(elavl3:shank3bΔC) transgenic fish (0.26 Hz), which express a dominant-negative isoform of shank3b, was lower than in wt siblings (0.35 Hz). Inset: mean swim velocity was indistinguishable between Dn(elavl3:shank3bΔC) (2.58 ± 0.15 cm/s) and wt siblings (2.63 ± 0.20 cm/s). C. Cumulative distribution of time engaged in swim events of different duration. During normal visuomotor coupling (dashed lines), shank3b^−/−^ fish spent 23.5% of the time swimming in comparison to 27.6% in wt siblings. During visuomotor perturbations (solid lines), swimming time increased to 47.8% in shank3b^−/−^ and to 51.6% in wt siblings. Inset: log ratio of swim time during perturbations over swim time during normal visuomotor coupling. D. Fraction of swim-modulated cells in the telencephalon of shank3b−/− fish was increased relative to wt siblings in Dm, cDc and Dl. E. Tail movement as a function of time and trials in shank3b−/− fish (right; n = 10) and wt siblings (left; n = 8). Trials were sorted by BREP onset. Trials below the yellow lines are negative trials (no BREP). F. Neuronal activity of pre-BREP neurons in Dm in shank3b−/− fish and wt siblings around the onset of a left-right reversal (white line). Activity of neurons in shank3b−/− fish was more heterogeneous.

Movie S1. Virtual reality and two-photon calcium imaging for adult zebrafish under head fixation. Related to Figure 1.

Movie S2. Behavioral responses to perturbations (BREPs). Related to Figure 3.

Movie S3. Post-surgery behaviors: saccades, locomotion, feeding and social affiliation. Related to STAR Methods.

